# *Enterocloster clostridioformis* induces host intestinal epithelial responses that protect against *Salmonella* infection

**DOI:** 10.1101/2023.07.20.549886

**Authors:** Benjamin S. Beresford-Jones, Satoshi Suyama, Simon Clare, Amelia Soderholm, Wangmingyu Xia, Puspendu Sardar, Katherine Harcourt, Trevor D. Lawley, Virginia A. Pedicord

## Abstract

Promoting resistance to enteric pathogen infection is a core function of the gut microbiota. However, many of the host-commensal interactions that likely mediate this protection remain uncharacterised. By screening gnotobiotic mice monocolonised with a range of mouse-derived commensal bacteria we have identified *Enterocloster clostridioformis* as a protective species against *Salmonella* Typhimurium infection. Unlike the colonisation resistance induced by some commensal bacteria, *E. clostridioformis* selectively induces a previously uncharacterised microbe adaptation response at the level of the caecal intestinal epithelium and the underlying mucosal immune system to mediate host-dependent resistance to infection. Triggering this pathway may therefore constitute a novel strategy to enhance protective responses against enteric infections.

## Introduction

Non-typhoidal salmonellosis (NTS) is a major global health concern, causing 93 million enteric infections and over 150,000 deaths each year^1, 2^. Furthermore, NTS results in 4.07 million disability-adjusted life years (DALYs), more than any other foodborne infection^3^. NTS is predominately acquired through consumption of contaminated animal products, especially cattle, swine, and poultry^4^, and drug resistant *Salmonellae* are now spreading in the human food supply chain increasing the financial burden of disease and worsening infection outcomes^5–7^. Effective treatment strategies and alternative approaches are therefore urgently required.

The gut microbiota has been implicated in providing colonisation resistance against *Salmonella* infection for nearly 70 years^8^. This colonisation resistance has been extensively studied, and many mechanisms have been identified by which commensal bacteria can inhibit colonisation of enteric pathogens: competition for metabolites including carbon sources, iron, oxygen, and electron acceptors^9–18^; production of inhibitory substances^19, 20^; and induction of host immune responses to indirectly control pathogen titres^21–23^.

However, the microbiota is also important for determining the pathophysiology of infection. Mice infected with *Salmonella* Typhimurium (*S.*Tm) do not develop robust gut inflammation akin to acute NTS, but instead develop a systemic, typhoid fever-like disease ^24, 25^. In contrast, antibiotic pre-treatment prior to *S.*Tm infection triggers an inflammatory diarrhoeal disease which mimics acute NTS in humans and can be used as a mouse model for *Salmonella* enterocolitis^8, 25–27^. This highlights the importance of microbiota-mediated mechanisms of protection that do not fall within the remit of colonisation resistance. For example, by modulating of *S.*Tm virulence^11, 28–31^ or inducing immune-mediated protection against infection and tolerance to immunopathology^32^, commensal bacteria can reduce the severity of disease without directly affecting the infectious titres in the intestine.

Although some mechanisms of commensal-induced host-mediated protection from enteric pathogens have been identified previously^33–36^, it is highly likely that many more examples and mechanisms of commensal-mediated enteric pathogen resistance remain unknown^32^. In this study, we have identified a novel function of commensal *Enterocloster clostridioformis* in the induction of protective responses in a mouse model of acute *Salmonella* enterocolitis.

We go on to show that *E. clostridioformis* localises in close proximity to the caecal epithelium and elicits both a unique antimicrobial response in caecal epithelial cells and a shift in local regulatory and effector T cell populations. These host effects correlated with decreased tissue pathology and significantly prolonged survival in germ-free mice. Collectively, these findings characterise a previously unknown role for *E. clostridioformis* in promoting host protection against infectious enterocolitis.

## Results

### An *in vivo screen* reveals *E. clostridioformis*-mediated protection against *Salmonella* infection

The gut microbiota has been known to play an important role in host protection against gastrointestinal infections since the seminal work by Margaret Bohnhoff and colleagues in 1952. We observed that different regimens of antibiotic treatment result in very different levels of susceptibility to infection with *S.*Tm strain 14028 in microbially replete (SPF) mice. Notably, germ-free mice were exquisitely sensitive to *S.*Tm infection, with pathogen titres of <100 CFU/mouse resulting in rapid weight loss and complete mortality within two days of infection (Figure 1a). Importantly, survival significantly correlated with the degree to which the microbiota remained intact (Figure 1b,c).

**Figure 1.**
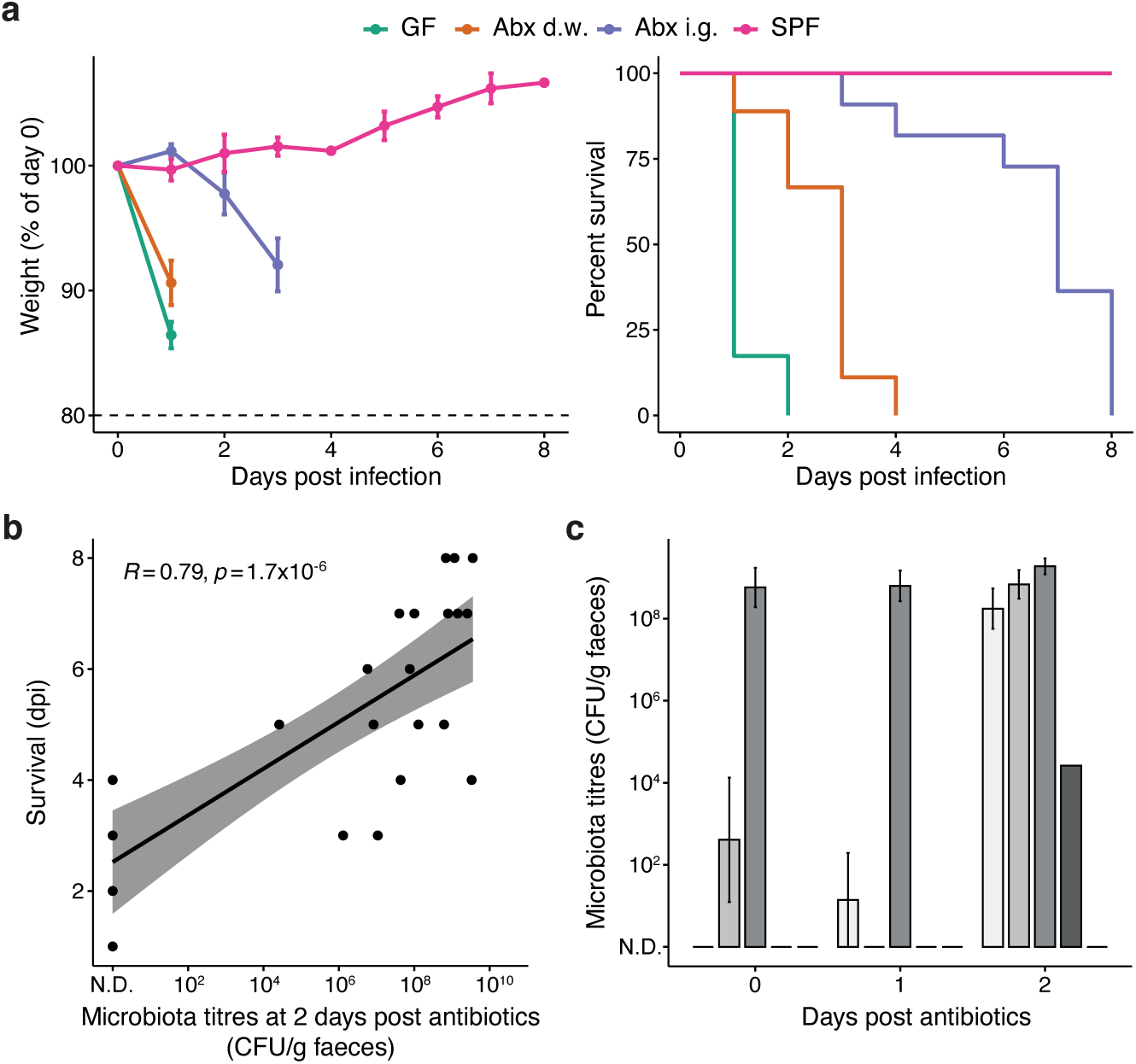
The gut microbiota is essential for resistance against Salmonella Typhimurium infection but can confound mouse studies. (a) Weight loss (left) and survival (right) data for C57BL/6NTac mice infected with *S.*Tm following different treatments: GF (n = 23); Abx d.w., AVMN antibiotics administered in the drinking water for 14 days (n = 9); Abx i.g., AVMN+Cf antibiotics administered by daily oral gavage for seven days (n = 11); SPF, specific pathogen-free (n = 3). *S.*Tm infectious doses: SPF = 10^6^ CFU/mouse; GF = 10^2^ CFU/mouse. (b) Faecal microbiota titres at two days post cessation of antibiotics correlate with duration of survival following *S.*Tm infection. Points represent individual mice. This subfigure combines data from both d.w. and i.g. protocols. Linear correlation was assessed using the Pearson product-moment correlation coefficient (PPMCC). Data for subfigures (a) and (b) are from three independent experiments. (c) Titres of faecal commensal bacteria in the days following antibiotic treatment in the drinking water. Each bar (different shades of grey) represents the mean faecal titres for a different cage of mice from a single experiment. Error bars indicate SEM.

To avoid the variance observed in antibiotic-treated animals, we therefore performed a screen of mouse-derived commensal microbes in germ-free mice to identify causative microbe-host interactions that are protective against *S.*Tm infection. Germ-free mice aged six to eight weeks old were monocolonised via intragastric administration with a single, high-titre inoculum of a commensal bacterial species isolate from the Mouse Culture Collection^37^. Following 14 days of monocolonisation, mice were orally infected with *S.*Tm 14028, and weight loss at 1 day post infection and time to reach 80% baseline weight were used as proxies for infection severity. A total of 18 commensal bacterial isolates from the Mouse Culture Collection were successfully screened for their impact on *S.*Tm infection (Table 1), including representatives from five taxonomic phyla (Figure 2a). No monocolonised mice suffered any mortality or exhibited any manifestations of disease during the pre-infection period.

**Figure 2.**
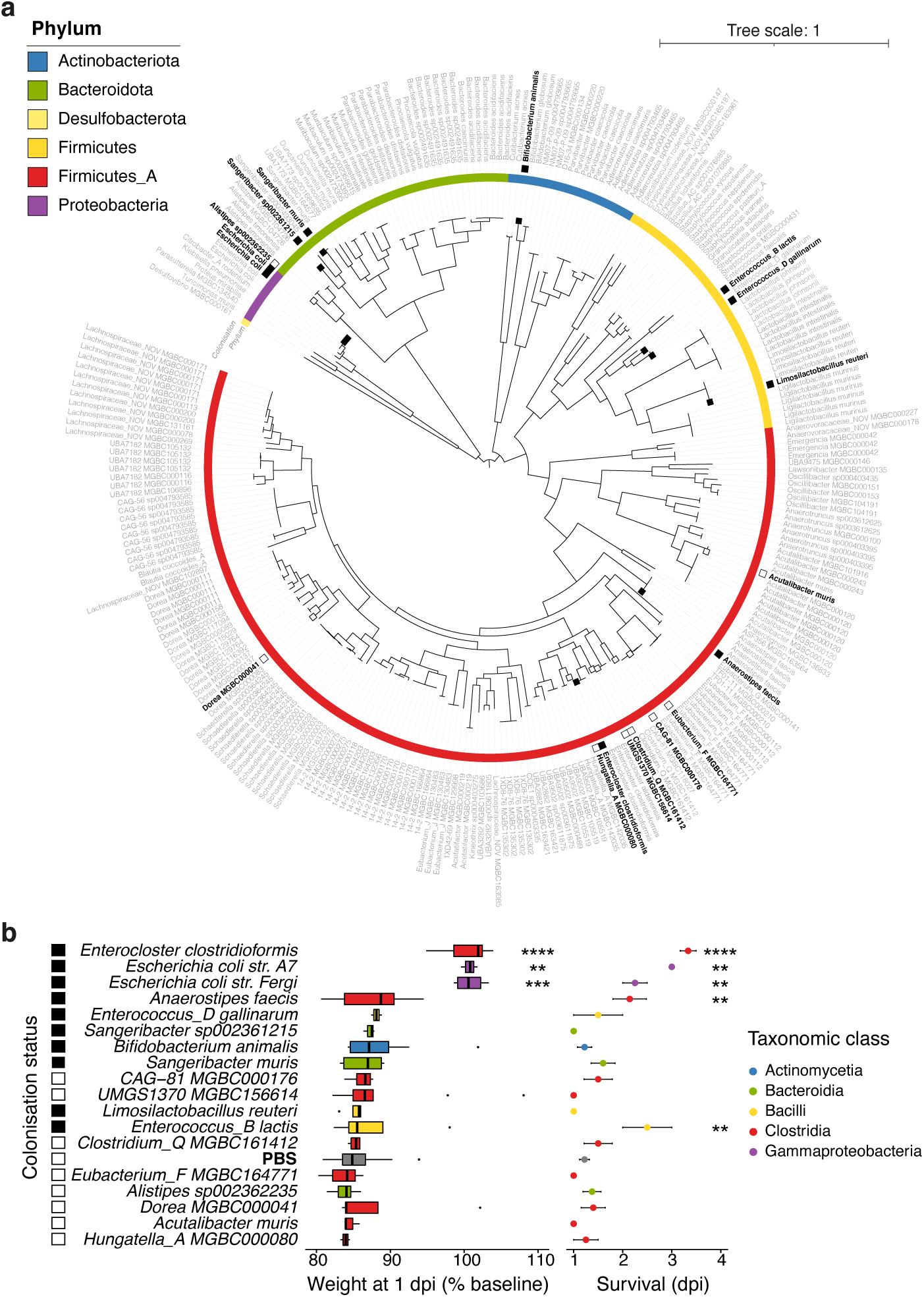
Screened isolates of the Mouse Culture Collection. (a) Maximum-likelihood phylogenetic tree of the 276 isolates of the MCC in addition to the genome of *Enterococcus_B lactis* str. Com15 (formerly *Enterococcus faecium*). Distances represent BLOSUM45 matrix analysis of amino acid alignments of 120 bacterial core genes using the JTT+CAT model. Screened isolates are indicated in black boldface type. Inner colour bar indicates the taxonomic phylum of isolates. Filled black boxes illustrate isolates that successfully colonised mice after 14 days monocolonisation, while filled white boxes indicate species that were administered to mice but could not be recovered from faeces after 14 days. (b) Monocolonisation with some commensals but not others induces resistance to *Salmonella* Typhimurium infection. From left to right: species name; filled boxes demarcate isolates that were recovered from the faeces 14 days after inoculation; mouse weights at 1 dpi as a percentage of starting weights (0 dpi); mouse survival post-infection. Statistical significance for both datasets was calculated with Kruskal-Wallis tests, using the PBS group as reference.

**Table 1.**
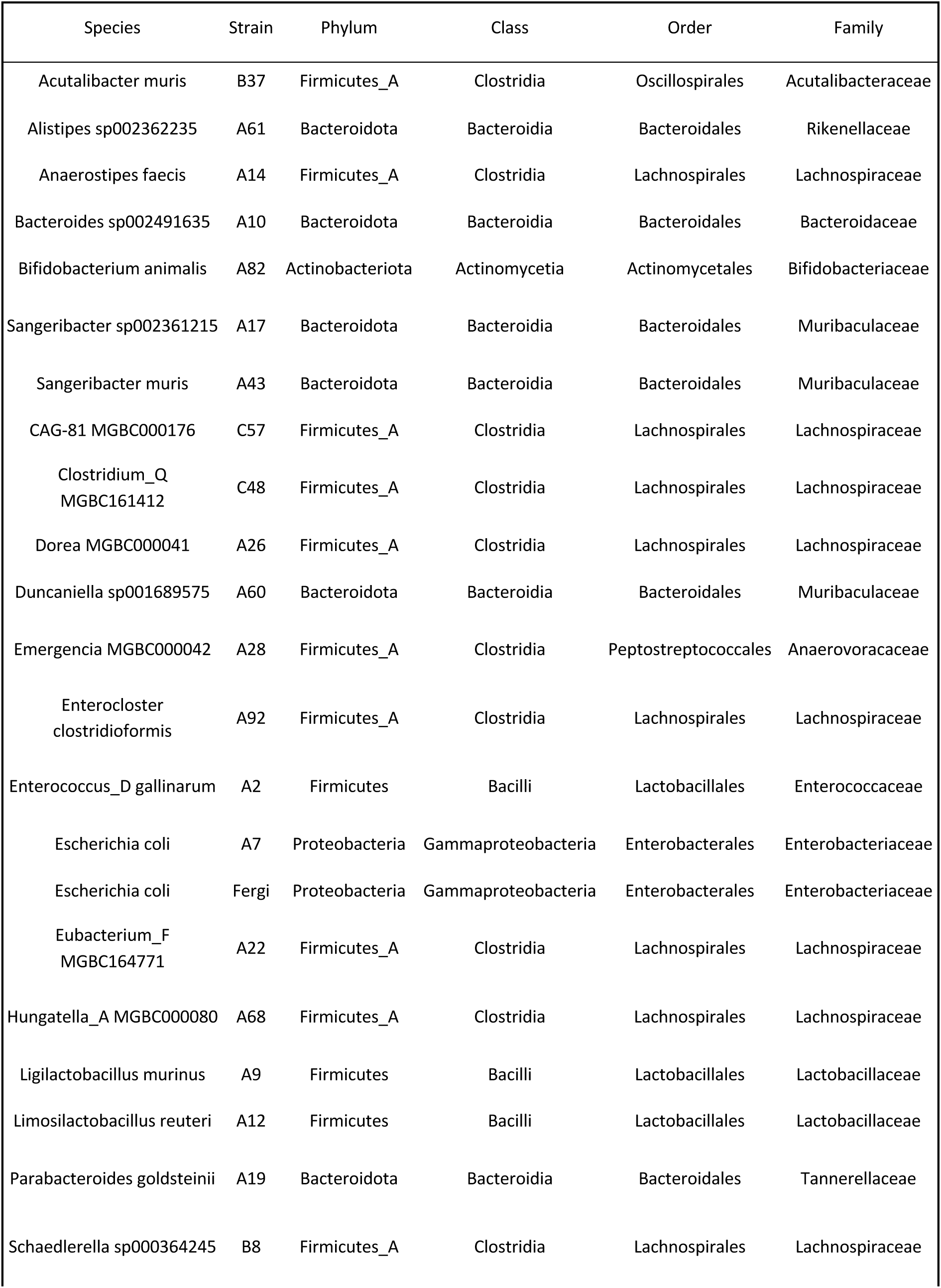
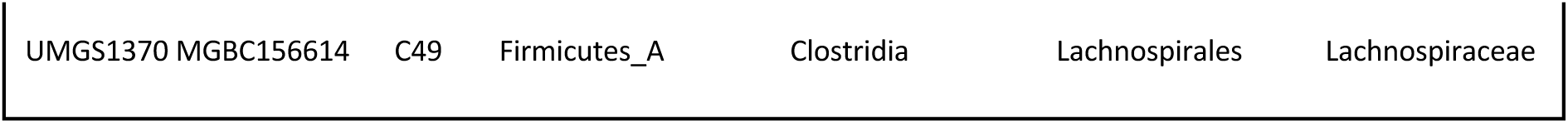
Isolates of the MCC used to screen for resistance against Salmonella Typhimurium infection.

| Species | Strain | Phylum | Class | Order | Family |
| --- | --- | --- | --- | --- | --- |
| Acutalibacter muris | B37 | Firmicutes_A | Clostridia | Oscillospirales | Acutalibacteraceae |
| Alistipes sp002362235 | A61 | Bacteroidota | Bacteroidia | Bacteroidales | Rikenellaceae |
| Anaerostipes faecis | A14 | Firmicutes_A | Clostridia | Lachnospirales | Lachnospiraceae |
| Bacteroides sp002491635 | A10 | Bacteroidota | Bacteroidia | Bacteroidales | Bacteroidaceae |
| Bifidobacterium animalis | A82 | Actinobacteriota | Actinomycetia | Actinomycetales | Bifidobacteriaceae |
| Sangeribacter sp002361215 | A17 | Bacteroidota | Bacteroidia | Bacteroidales | Muribaculaceae |
| Sangeribacter muris | A43 | Bacteroidota | Bacteroidia | Bacteroidales | Muribaculaceae |
| CAG-81 MGBC000176 | C57 | Firmicutes_A | Clostridia | Lachnospirales | Lachnospiraceae |
| Clostridium_Q<br>MGBC161412 | C48 | Firmicutes_A | Clostridia | Lachnospirales | Lachnospiraceae |
| Dorea MGBC000041 | A26 | Firmicutes_A | Clostridia | Lachnospirales | Lachnospiraceae |
| Duncaniella sp001689575 | A60 | Bacteroidota | Bacteroidia | Bacteroidales | Muribaculaceae |
| Emergencia MGBC000042 | A28 | Firmicutes_A | Clostridia | Peptostreptococcales | Anaerovoracaceae |
| Enterocloster<br>clostridioformis | A92 | Firmicutes_A | Clostridia | Lachnospirales | Lachnospiraceae |
| Enterococcus_D gallinarum | A2 | Firmicutes | Bacilli | Lactobacillales | Enterococcaceae |
| Escherichia coli | A7 | Proteobacteria | Gammaproteobacteria | Enterobacterales | Enterobacteriaceae |
| Escherichia coli | Fergi | Proteobacteria | Gammaproteobacteria | Enterobacterales | Enterobacteriaceae |
| Eubacterium_F<br>MGBC164771 | A22 | Firmicutes_A | Clostridia | Lachnospirales | Lachnospiraceae |
| Hungatella_A MGBC000080 | A68 | Firmicutes_A | Clostridia | Lachnospirales | Lachnospiraceae |
| Ligilactobacillus murinus | A9 | Firmicutes | Bacilli | Lactobacillales | Lactobacillaceae |
| Limosilactobacillus reuteri | A12 | Firmicutes | Bacilli | Lactobacillales | Lactobacillaceae |
| Parabacteroides goldsteinii | A19 | Bacteroidota | Bacteroidia | Bacteroidales | Tannerellaceae |
| Schaedlerella sp000364245 | B8 | Firmicutes_A | Clostridia | Lachnospirales | Lachnospiraceae |

Although all isolates were successfully cultured from the oral gavage inoculum, only ten isolates were viably recovered from the faeces after 14 days of monocolonisation. Most of these non-recoverable species were members of the class Clostridia (n = 7; phylum Firmicutes_A), while one member of the class Bacteroidia, *Alistipes sp002362235*, was also not recovered. Successful colonisation with any isolate tended to reduce the weight loss at 1 day post-infection, as PBS controls lost more weight than any successfully monocolonised mice on average. *E. clostridioformis* and two *Escherichia coli* strains, A7 and Fergi, prevented weight loss at 1 day post infection and significantly increased survival compared to PBS controls (Figure 2b). Two additional isolates, *Anaerostipes faecis* and *Enterococcus_B lactis*, also significantly improved mouse survival compared to PBS controls, but did not reduce weight loss at 1 day post-infection.

### *E. clostridioformis* elicits tissue protection without preventing *Salmonella* colonisation

The *Enterobacteriaceae* family, as well as individual strains of *E. coli* (e.g., Nissle 1917), have previously been associated with protection against S.Tm infection in mice through mediating colonisation resistance. Indeed, *E. coli* has become a model organism for understanding colonisation resistance in *S.*Tm infection^38^. In contrast, *E. clostridioformis* has not been previously implicated in protection against *S.*Tm infection. Therefore,we sought to better understand this association. Although all mice did eventually proceed to mortality, monocolonisation with *E. clostridioformis* significantly increased resistance to *S.*Tm infection, delaying and reducing the rate of weight loss (Figure 3a) and increasing survival (Figure 3b). Mortality at 1 day post-infection was reduced from 76% in germ-free mice to 0% following *E. clostridioformis* treatment.

**Figure 3.**
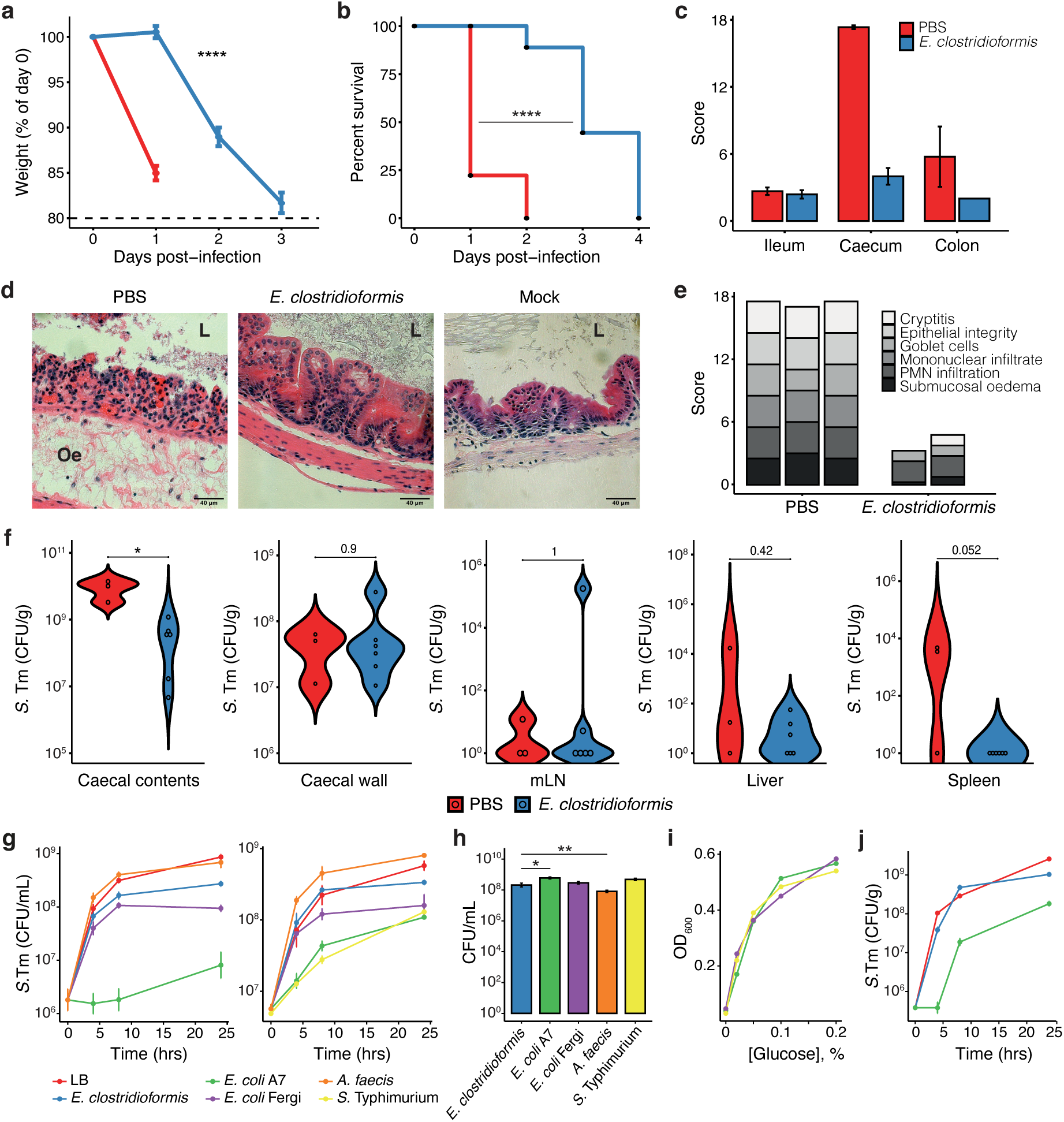
*Enterocloster clostridioformis* protects germ-free mice against *Salmonella* Typhimurium infection independently of colonisation resistance. (a) Weight loss and (b) survival following infection with 10^2^ CFU/mouse S.Tm. Weight loss data represent the mean ± SEM; data for timepoints after a mouse was euthanised are censored. Kruskal-Wallis and log-rank tests were used for statistical analyses in (a) and (b) respectively; for survival, p = 5×10^−9^. Data represent 36 mice across four experiments. (c) Cumulative pathology scores for H&E-stained sections of the ileum, caecum, and colon of GF or E. clostridioformis-monocolonised mice at 1 dpi. Data represent mean ± SEM of cumulative pathology scores from individual mice; PBS, n = 3; E. clostridioformis, n = 2. (d) Representative sections of the caecum at 1 dpi (PBS, *E. clostridioformis*) or from non-infected GF mice (Mock). L = caecal lumen; Oe = submucosal oedema. Scale bars indicate 40 μm. (e) Breakdown of the caecum pathology scores at 1 dpi according to the six contributing subcomponents. Each bar represents an individual mouse. (f) S.Tm titres (CFU/g tissue) at 1 dpi in the caecum luminal contents, caecal wall, mLN, liver, and spleen (left to right). Each data point represents a single mouse; Mann-Whitney U tests were used to assess statistical significance. Subfigures (c–f) represent data from a single experiment. (g) Growth titres of *S.*Tm in commensal isolate stationary phase broth (left) and spent media (right) over time. (h) Commensal and *S.*Tm titres at 24 hours in LB broth under anaerobic conditions. (i) Growth response of *S.*Tm to glucose supplementation of sterile-filtered spent LB broth. (j) Growth of *S.*Tm when spiked into faeces from GF or monocolonised mice.

*S.*Tm produces virulence factors which it both secretes and directly injects into host intestinal epithelial cells (IECs) via its SPI-1-encoded type III secretion system to mediate invasion, resulting in an acute inflammatory response. Accordingly, at 1 day post-infection germ-free mice exhibited extensive pathological signs in the caecum (Figure 3c), with extreme submucosal oedema, complete loss of epithelial integrity, and goblet cell depletion (Figure 3d,e). In comparison, *E. clostridioformis*-monocolonised mice displayed markedly reduced *S.*Tm-induced caecal inflammation and pathology at 1 day post-infection (Figures 3c-e). Although some pathological changes were seen in the colons of germ-free mice following *S.*Tm infection, most notably submucosal oedema, these were not as profound as in the caecum, and no mice from either cohort exhibited extensive pathology in the terminal ileum. This aligns with the previously reported invasion and pathology in the caecum and colon at early time points after *S.*Tm infection^26^.

Following on from the histological phenotype, we next considered the extent of intestinal and infection versus dissemination to peripheral organs. At 1 day post-infection, the S.Tm titres in the caecal lumen were significantly lower in *E. clostridioformis*-monocolonised mice than in germ-free controls (Figure 3f); however, they were still high enough to mediate invasion, as titres of *S.*Tm in the caecum wall were equivalent between these cohorts. There were no differences in mesenteric lymph node (mLN) or liver titres between the cohorts, although there was a trend of reduced systemic infection as indicated by lower *S.*Tm titres in the spleen (p = 0.052).

The best described mechanism for commensal-mediated protection against *S.*Tm is colonisation resistance^38^. Indeed, *E. coli* is well-known for its capacity to protect against *S.*Tm through mechanisms of colonisation resistance^18, 39, 40^, and it is therefore unsurprising that *E.* coli isolates ranked among the topmost protective species from our screen. As the dynamics of *E. clostridioformis*-mediated protection are similar to those of the *E. coli* strains, we assessed whether this species also mediated resistance to infection by colonisation resistance. To separate inter-bacterial dynamics from the host context, we performed a series of *in vitro* and *ex vivo* competition assays to query whether *E. clostridioformis* can directly inhibit or compete with *S.*Tm. We found that *S.*Tm growth was significantly attenuated when grown in stationary phase broth culture of *E. coli* A7, but not *E. clostridioformis* or *A. faecis*, and a similar trend was observed for *S.*Tm growth in sterile-filtered spent media, where spent media of *E. coli* A7 was equivalent to spent media from *S.*Tm itself at attenuating pathogen growth (Figure 3g). In contrast, the growth of *S.*Tm in stationary or spent media of *E. clostridioformis* was similar to its growth in LB, although *S.*Tm titres at 24 hours were significantly reduced in the *E. clostridioformis* spent media, likely due to a partial overlap in metabolic nutrient utilisation.

While the commensal cultures used for these assays exhibited some variation in growth, all differences were less than 10-fold (1 log) and were therefore unlikely to be biologically significant (Figure 3h). The reduced stationary phase titres of *S.*Tm in spent media from the *E. coli* strains and *S.*Tm itself were mediated by carbon source depletion rather than media toxification, as addition of glucose to the spent media supplemented *S.*Tm growth in a concentration-dependent manner (Figure 3i). Lastly, we performed *ex vivo* faecal spiking assays to confirm the absence of a colonisation resistance phenotype in a more physiologically relevant setting. The faeces of mice monocolonised with either *E. coli* A7 or *E. clostridioformis* were spiked with *S.*Tm. While *E. coli* A7 monocolonisation significantly reduced and delayed *S.*Tm growth, *S.*Tm growth in *E. clostridioformis*-monocolonised faeces was equivalent to growth in germ-free faeces (Figure 3j). Taken together, these findings strongly suggest that the protective phenotype of *E. clostridioformis* against *S.*Tm is not mediated by colonisation resistance and requires host factors.

### *E. clostridioformis* colonisation induces host antimicrobial responses in the caecal epithelium

To examine the effects of these gut commensal bacteria on the host we investigated whether monocolonisation with *E. clostridioformis* was associated with changes in gene expression in the intestinal epithelium. We performed RT-qPCR on IECs from the small intestine, caecum, and colon from germ-free and microbiota-replete SPF mice, as well as mice monocolonised with *E. clostridioformis, E. coli* A7 or *A. faecis*. We observed monocolonisation-induced changes in IEC gene expression that were both intestinal region– and commensal microbe-specific. Most notably, monocolonisation with *E. clostridioformis* induced a six-fold increase in caecal IEC expression of the antimicrobial peptide resistin-like molecule b (RELMb, gene: *Retnlb*; Figure 4a). As *Retnlb* expression was not increased following monocolonisation with *E. coli* or *A. faecis*, this response was specific to *E. clostridioformis* and did not reflect a general response to the presence of bacterial colonisation. Furthermore, *E. clostridioformis* specifically upregulated this pathway, as monocolonisation was not associated with increased expression of any of the other barrier function genes we examined, including *Reg3g*, *Fut2*, and *Muc2* (Figure 4b). This response was also anatomically restricted as *E. clostridioformis* monocolonisation induced *Retnlb* expression to a lesser extent in the colon and not at all in the IECs of the terminal ileum.

**Figure 4.**
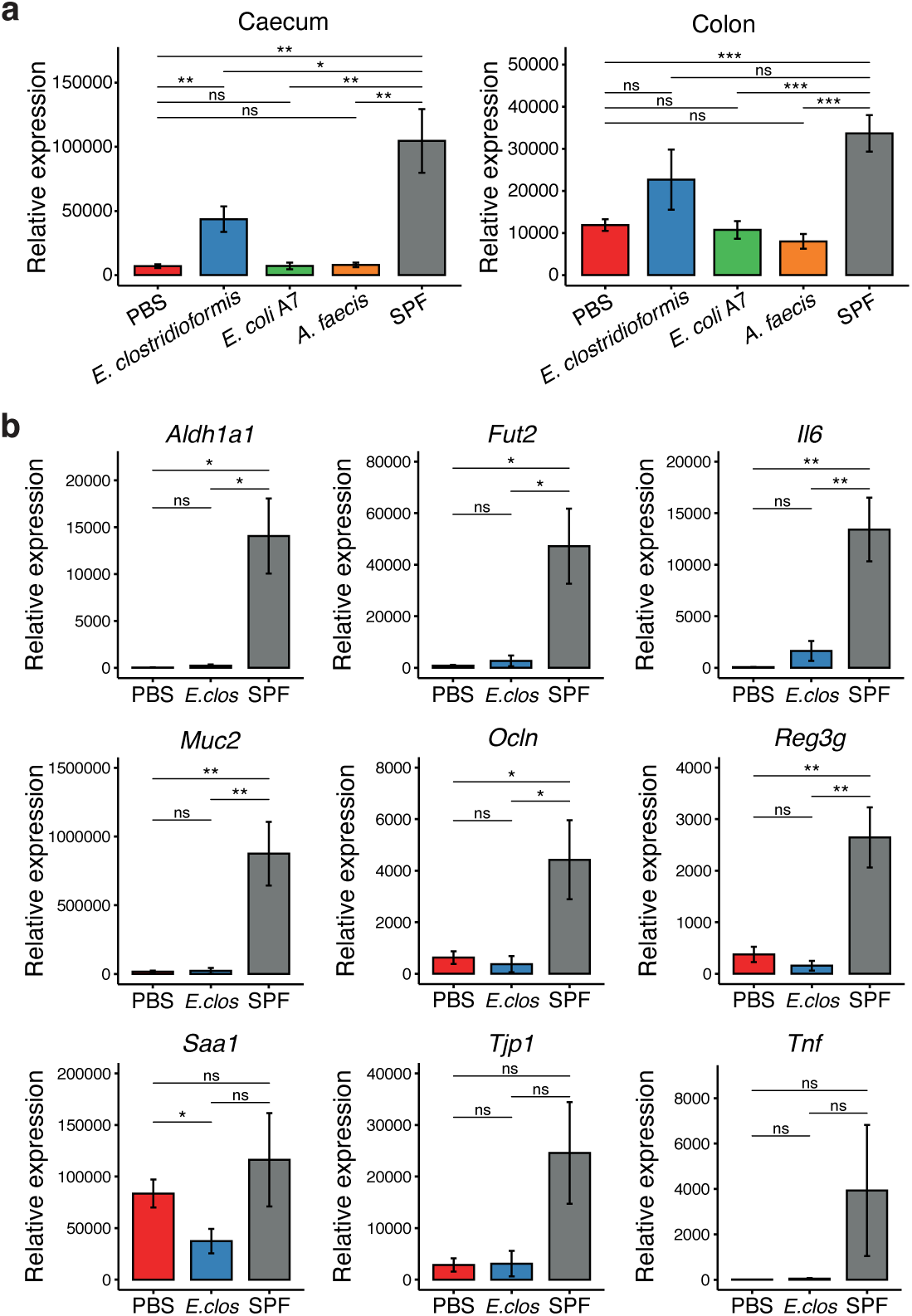
Enterocloster clostridioformis induces expression of Retnlb by the caecal epithelium. (a) Caecal (left) and colon (right) intestinal epithelial cell (IEC) *Retnlb* expression from GF, commensal-monocolonised, or SPF mice. (b) Gene expression profiles of caecal IECs from GF, *E. clostridioformis*-monocolonised or conventional SPF mice. Student’s t-tests were used for statistical comparisons.

As gene expression changes can be induced by bacteria via a range of mechanisms, including physical contact with host tissues or through secretion of proteins or small molecules, we performed fluorescence in situ hybridisation (FISH) with a universal 16S rRNA gene probe to visualise commensal microbes in the context of the intestinal epithelium. Use of a modified mucus penetration score indicated that *E. clostridioformis* was significantly more closely associated with the caecal epithelium when compared to *E. coli* A7, *A. faecis*, and the multispecies microbiota of SPF mice (Figure 5a). This trend did not continue into the colon where mucus penetration was equivalent between these bacterial species (Figure 5b). Of note, SPF mice exhibited a more structured inner mucus layer in both the caecum and the colon (Figure 5c). Some bacterial species that make physical contact with the intestinal epithelium are known to induce changes in gene expression, most notably SFB and SAA1/2. Although *E. clostridioformis* appeared to be more closely associated with the caecal epithelium than *E. coli* A7 and *A. faecis*, expression of *Saa1* was significantly reduced following *E. clostridioformis* monocolonisation than compared to germ-free mice (Figure 4b).

**Figure 5.**
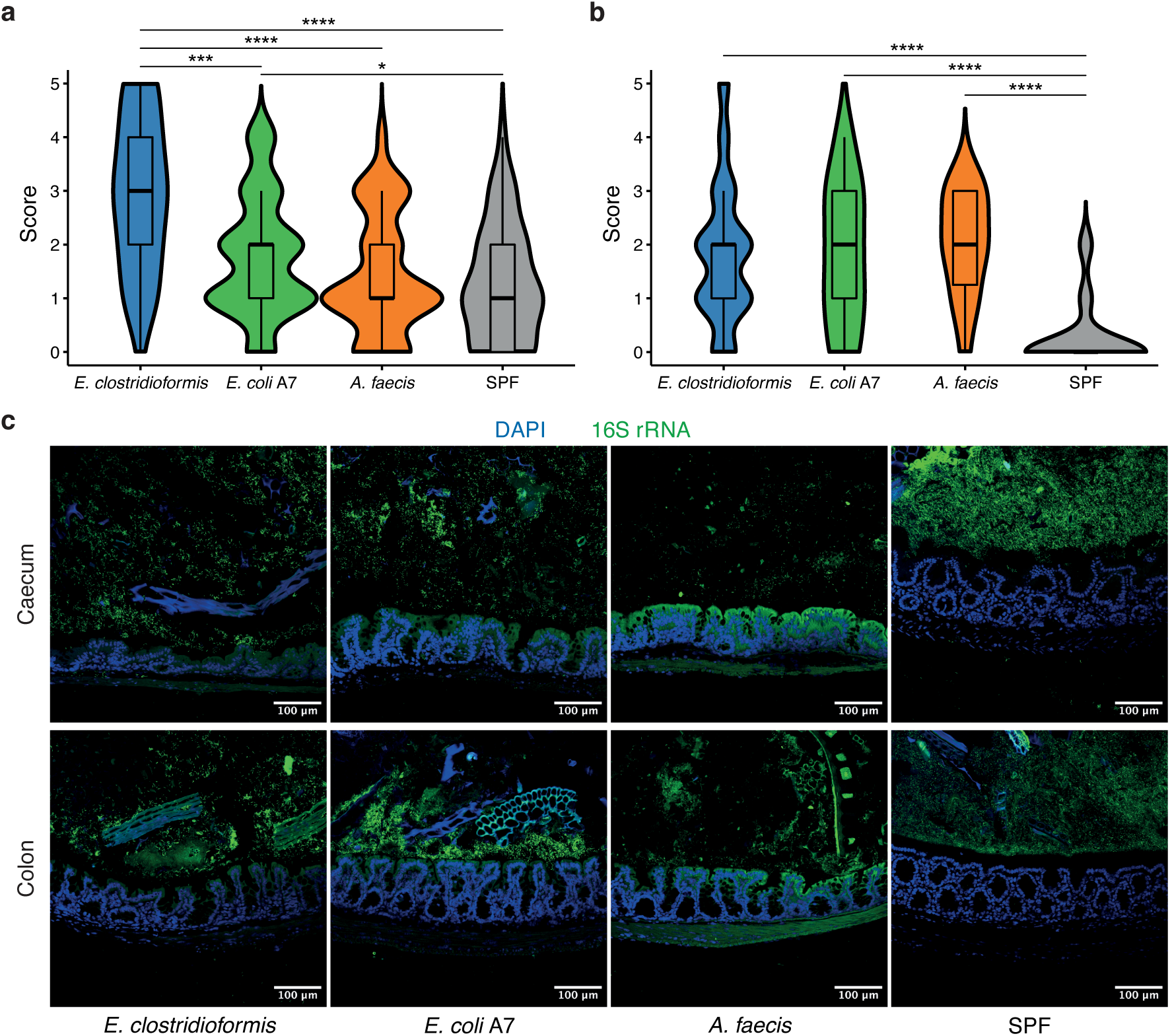
Enterocloster clostridioformis is closely associated with the caecal epithelium. (a–b) Mucus penetration scores in the caecum (a) and colon (b) for *E. clostridioformis*, *E. coli* A7, or *A. faecis* monocolonised GF mice, as well as control SPF mice. Scoring was performed by an independent and blinded investigator. (c) Representative micrographs of microbes in the context of the caecal (upper row) and colonic (lower row) epithelium. Nuclei are stained with DAPI (blue), and bacteria are stained with a universal 16S rRNA FISH probe conjugated to FITC (green). Scale bars indicate 100 μm.

To assess how these changes at the epithelial barrier affect *S.*Tm pathology and invasion, we next examined the caecal epithelium of monocolonised mice at 1 day post-infection. *S.*Tm was abundant in the caecal lumens of mice from both treatment groups although more *S.*Tm could be observed in germ-free mice than their *E. clostridioformis*-monocolonised counterparts, in keeping with the increased luminal titres observed in Figure 3f. The distribution of *S.*Tm also differed between treatment groups, with larger clumps of *S.*Tm found close to the epithelium in germ-free mice and more evenly distributed pathogen in the lumens of *E. clostridioformis*-monocolonised mice (Figure 6a). *S.*Tm extensively effaced the epithelium in germ-free mice but exhibited much less contact with the caecal epithelium in *E. clostridioformis*-monocolonised mice.

**Figure 6.**
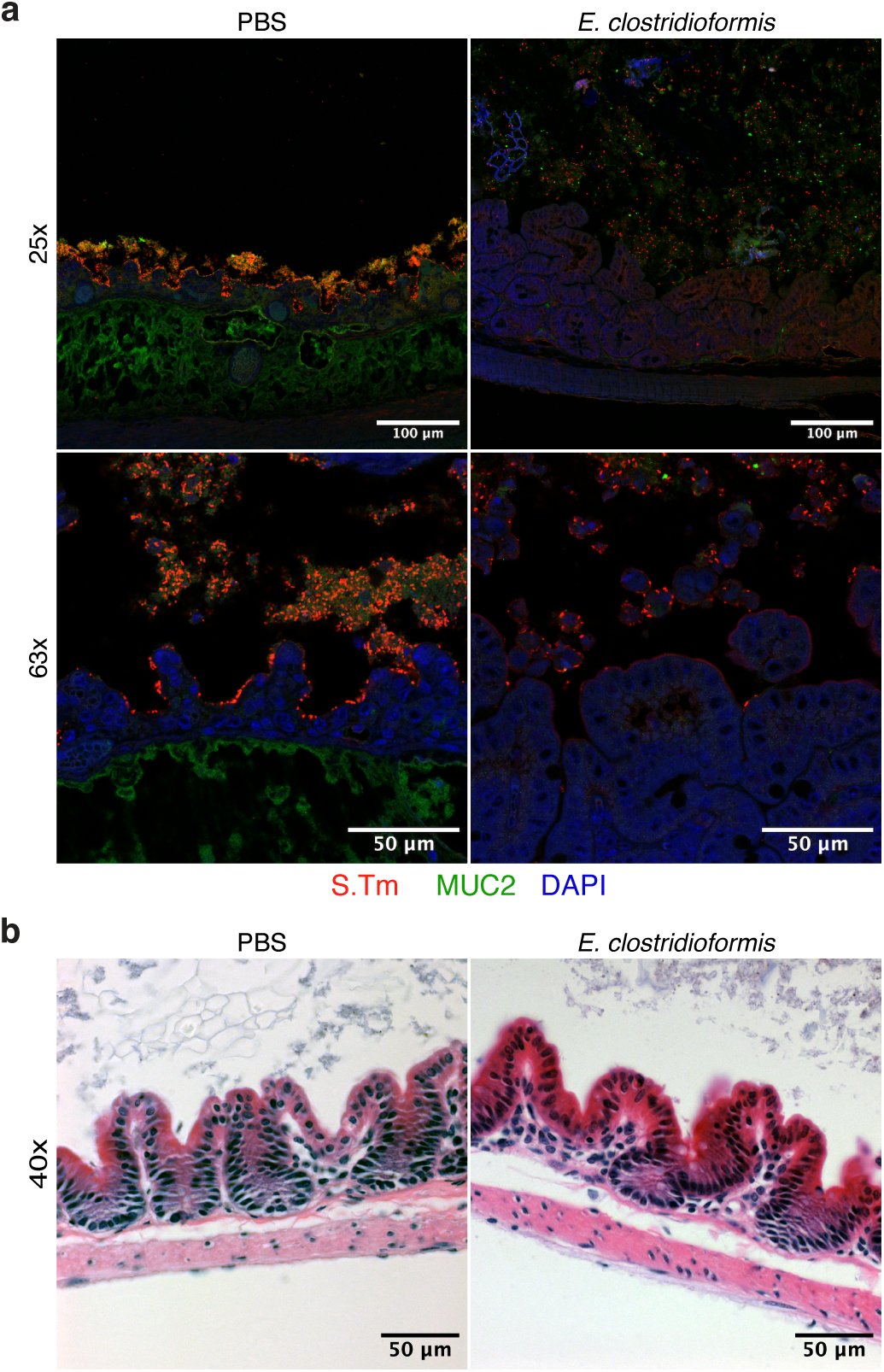
E. clostridioformis monocolonisation is associated with a hyperproliferative epithelial response and less pathogen–epithelium contact. (a) Immunofluorescence of *S.*Tm (red), MUC2 (green), and host cells (blue) at 1 dpi in mock-treated GF mice (‘PBS’, left column panels) and *E. clostridioformis*-monocolonised mice (right column panels). Submucosal oedema appears green, most likely due to incomplete blocking of endogenous mouse antibodies. Figures are representative of findings from a single experiment with three PBS and two *E. clostridioformis* mice. (b) H&E histology sections of mock-treated and *E. clostridioformis*-monocolonised mice prior to infection. Mice had been monocolonised for 14 days. Figures are representative of findings from three experiments. Scale bars are provided in the bottom right of each panel.

*E. clostridioformis* monocolonisation was associated with a thickened epithelium with deeper crypts and the development of small, villi-like extensions into the lumen, together indicating a hyperproliferative response of the epithelium to infection (Figure 6a). This proliferative phenotype was specific to the post-infection context, as no such differences could be observed between the epithelia of germ-free or *E. clostridioformis*-monocolonised mice prior to infection (Figure 6b). As *S.*Tm makes contact with the epithelium in both treatment groups, but less so following *E. clostridioformis* monocolonisation, this proliferative phenotype may represent a microbiota-enabled response to infection resulting in more effective barrier function and leading to a subsequent protective phenotype.

### Anti-inflammatory T cell ratios are increased by *E. clostridioformis*

Finally, we considered the impact of commensal monocolonisation on the mucosal immune system. The gastrointestinal tract is the largest immune organ in the body, with more lymphocytes than the spleen. These lymphocytes are distributed throughout different compartments of the intestines, with each compartment hosting different subsets of immune cells. Using flow cytometry of intestinal cells isolated from germ-free or monocolonised mice, we quantified the relative abundance of lymphocyte populations in the intraepithelial compartments and lamina propria of the small intestine, caecum, and colon, in addition to the Peyer’s patches and mLNs. Monocolonisation with both *E. clostridioformis* and *E. coli* A7 induced a significant increase in the relative abundance of CD4+ T cells in the caecal lamina propria lymphocyte (LPL) (Figure 7a). However, the nature of the CD4+ T cell expansion was specific to each commensal isolate. *E. clostridioformis* induced a selective increase in FoxP3+ CD4+ Tregs, resulting in an increased Treg:Teffector cell ratio. In contrast, *E. coli* A7 colonisation induced the expansion of CD4+ T effector cells, significantly reducing the Treg:T effector cell ratio. *E. clostridioformis* colonisation also increased the relative abundance of Tregs in the colonic LPL compartment (Figure 7b) as well as increased the Treg:Teffector cell ratios in the intraepithelial lymphocyte (IEL) compartments of the caecum, colon, and small intestine (Figures 7c). There was no difference in Treg cell proportions in mLNs, Peyer’s patches, or in the small intestinal LPL compartment. The proportion of CD8ab+ T cells was unaffected by *E. clostridioformis* monocolonisation, as were other IEL populations, including CD8aa+, CD8aa+ CD8ab+, and double negative (CD4– CD8–) T cells. These findings may represent a novel mechanism of microbiota-induced host protection against the infection-associated inflammation and tissue damage.

**Figure 7.**
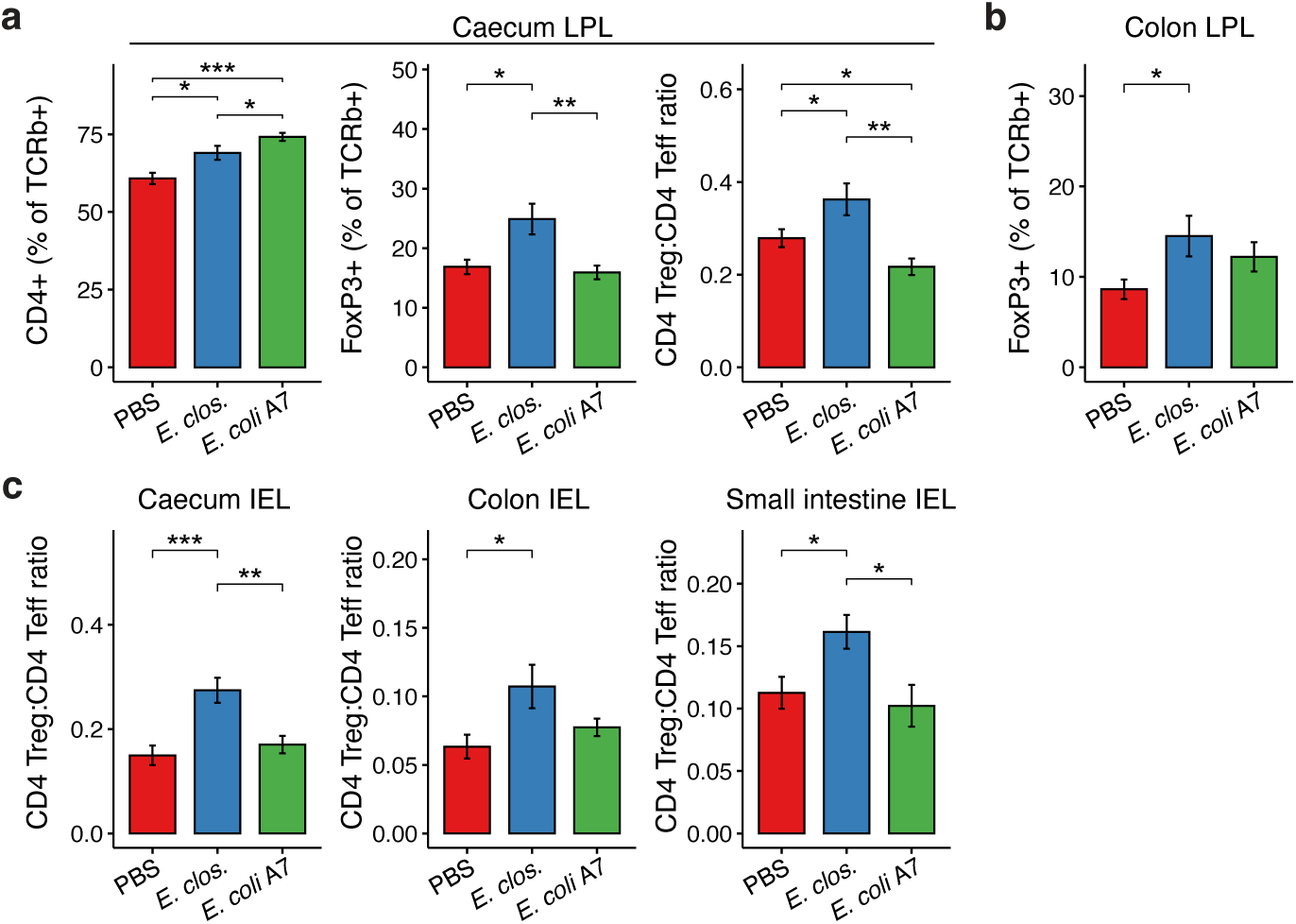
Enterocloster clostridioformis monocolonisation increases the proportion of Tregs in the caecal epithelium. Flow cytometry was used to quantify gut lymphocyte compartments. Single cells were pre-gated using forward scatter, side scatter, and cell width, and Viable (Live/Dead Aqua−) CD45+ cells were then gated for analyses. T cells were gated as CD3ε+ TCRβ+. (a) Caecal LPL compartment: CD4+ T cells as a percentage of all αβ-T cells (left); FoxP3+ Tregs as a percentage of αβ-T cells (middle); ratio of CD4+ FoxP3+ Tregs to CD4+ FoxP3−T effector cells (right). (b) FoxP3+ Tregs as a percentage of αβ-T cells in the colonic lamina propria. (c) Ratio of CD4+ FoxP3+ Tregs to CD4+ FoxP3−T effector cells in the intraepithelial compartments of the caecum (left), colon (middle), and small intestine (right). Figures represent data from three independent experiments. Statistical significance was assessed using Wilcoxon tests.

## Discussion

In this paper, we utilised mouse-derived gut bacterial isolates to screen for microbial species that are associated with resistance to *Salmonella* infection in a germ-free mouse model. Through this work, we identified *E. clostridioformis* as a novel protective species against acute salmonellosis. Mice monocolonised with *E. clostridioformis* exhibited reduced weight loss and significantly prolonged survival with a concomitant reduction in caecal and colonic pathology. This phenotype was associated with increased expression of the AMP *Retnlb* in epithelial cells and a relative increase in CD4+ Tregs in the intraepithelial compartment and lamina propria. These findings were not general responses to bacterial colonisation but were instead specific to colonisation with *E. clostridioformis*.

This is the first study to associate *E. clostridioformis* with outcomes of *S.*Tm infection; however, a recent study has demonstrated a role for this species in *E. coli*-mediated colonisation resistance to *S.*Tm^40^. This study found that *E. clostridioformis* and *S.*Tm share complete overlap in metabolic capacity for C5 and C6 sugar sources, explaining why *E. clostridioformis* competes with *S.*Tm in this niche, and this is in keeping with our finding that monocolonisation with *E. clostridioformis* does significantly reduce caecal titres of *S.*Tm. However, by monocolonising mice with *E. clostridioformis*, we have shown that *E. clostridioformis*-mediated protection does not require *E. coli* or the colonisation resistance it elicits. In addition, several of our results indicate that the protective phenotype induced by *E. clostridioformis* monocolonisation is not mediated by colonisation resistance. Firstly, the extent of protection achieved by *E. coli* is slightly less than *E. clostridioformis*, despite exhibiting a much stronger colonisation resistance phenotype against *S.*Tm. Secondly, the infectious titres identified in the mural compartment at 1 day post-infection are equivalent between germ-free and *E. clostridioformis* monocolonised mice, indicating that titres in the caecal lumen are still sufficient for tissue invasion^26^. *E. clostridioformis* has also been previously associated with protection against *Clostridioides difficile* infection^41^. Although *C. difficile* is another gastrointestinal pathogen, it causes pathology through toxin production in contrast to the enteroinvasive mechanisms of *S.*Tm. Enteroinvasive and toxigenic mechanisms of pathogenesis are very distinct processes, and this further suggests that a common protective mechanism induced by *E. clostridioformis* is likely to occur at the level of the host instead of through direct interbacterial colonisation resistance.

SPF mice are commonly used models for the study of *S.*Tm infection. Although they represent a more physiologically replete system compared to germ-free mice, their use is associated with known caveats when applied to studying host-commensal interactions. Variation in the underlying microbiota of SPF mice can confound studies and hinder accurate observation and reproduction of microbial associations. We have shown that the speed of recovery of the microbiota following antibiotic treatment varies significantly between cages with implications for the response to infection, which correlated with both number and nature of bacterial growth following antibiotic therapy. Presence and abundance of the microbe of interest in the endogenous gut microbiota can confound experiments^42^, and would need to be established for each sample for SPF models to be used reliably. While some research groups embrace this endogenous variation between the microbiota of SPF mice to identify microbial factors that associate with phenotypes of pathogen resistance^18, 32^, these limitations indicate that SPF models are likely not sensitive nor reproducible enough for screening for protective host-commensal interactions against *S.*Tm infection. When considered in the context of unnecessary levels of mortality associated with antibiotic treatment protocols, germ-free mice become the model of choice for causative host-commensal microbiota studies.

Monocolonisation of germ-free mice induces host adaptation to commensal microbial signals, and therefore can determine how these responses are important in resistance to enteric pathogen infection. This cannot be easily assessed using SPF mice, which lend themselves more to modelling colonisation resistance. Additionally, the germ-free model allows commensal microbes to be associated with the host for prolonged periods of time. This is not reliably the case in SPF mice, as colonising microbes may be outcompeted by the resident microbiota or killed by incomplete clearance of administered antibiotics. Although some studies use repeated administration of microbes to SPF mice to prolong exposure of the host to a commensal species^43^, there is limited evidence to suggest that this improves colonisation or that it reliably exposes the host to commensal antigens or metabolites. We chose a monocolonisation period of two weeks for our screen. This length of time reduces the need for cage changes and therefore the rate of contamination, but also facilitates adaptation and equilibration of the host to microbial colonisation^44^. Furthermore, this length of time would suffice to establish whether monocolonisation with any commensal species in and of itself would cause acute or subacute pathology in the host. Some microbiota-dependent processes do require longer than two weeks. One such example is normalisation of the colonic mucus layer, which can take up to six weeks following reconstitution of the gut microbiota^45^. Phenotypes dependent on such processes would therefore not have been identified as part of our screen.

While we have not established a causal link between RELMb expression or Treg induction and *E. clostridioformis*-induced resistance to *S.*Tm, both represent promising mechanisms for further study. Tregs are known to be important in directing early adaptive immune responses to S.Tm infection^46^, and are also reduced in germ-free mice^47–49^. An increase in their relative proportion could represent a mechanism by which *E. clostridioformis* mediates tolerance to acute infection through a reduction in overt immune-mediated damage to host tissues^50^. Butyrate synthesis is an established mechanism by which commensal microbes can increase LPL Treg populations, however *E. clostridioformis* does not encode either terminal pathway required for producing butyrate implying that it utilises a different mechanism. Interestingly, this species has previously been identified in a screen for commensals that can induce Tregs proliferation^51^ although it was not followed up by this study and as such its Treg-inducing mechanism remains unknown. As our data indicate that *E. clostridioformis* monocolonisation increases the caecal Treg proportion of total CD4+ T cells to 37% – similar to published values for colonic LPLs in SPF mice^48, 52^ – further study of this microbe-host interaction is warranted beyond to its *S.*Tm resistance phenotype.

Similarly, induction of RELMb could represent a mechanism by which *E. clostridioformis* mediates its protective effects. While it has not previously been considered in the context of *S.*Tm, it is known to be directly bactericidal against Gammaproteobacteria such as *Citrobacter rodentium* and has a role in promoting microbiota spatial segregation in the colon^53^. Furthermore, RELMb is implicated in regulation of intestinal barrier function^54^ and has been previously associated with inducing a T cell-dependent IEC proliferation phenotype that is protective against enteropathogenic infection with *C. rodentium*^55^. It remains to be seen through further studies which of the potential Treg and RELMb mechanisms, if either, are important to the protective phenotype against *S.*Tm or whether they represent different elements of the same process.

NTS represents an important global health concern, and rising antibiotic resistance necessitates the investigation of alternative approaches to disease prevention and treatment. Our study provides the basis for further exploration of *E. clostridioformis* as a mediator of resistance to *S.*Tm infection and expands the current understanding of how the host adapts to the gut microbiota, highlighting novel pathways and mechanisms that could potentially be harnessed for prevention and treatment of NTS, as well as other pathologies.

## Materials and Methods

### Mice

Mice were maintained under either specific pathogen-free (SPF) or germ-free (GF) conditions at the Home Office-approved Wellcome Sanger Institute mouse facility and all procedures carried out in accordance with the United Kingdom Animals (Scientific Procedures) Act of 1986.

SPF mice were bred and maintained under barrier conditions in individually ventilated cages (IVCs), which were then transferred from the barrier to a separate containment room prior to experiments to facilitate researcher access. Male and female mice were housed separately and only female C57BL/6NTac mice aged six to eight weeks were used for SPF infection experiments. Non-acidified autoclaved water and irradiated chow (Safe R03-10 complete breeding diet; crude protein: 21.4%, crude fat: 5.1%, crude ash: 5.4%, crude fibre: 4.0%) were provided for *ad libitum* consumption. All cages were provided with bedding, nesting material, and enrichment. Mice were exposed to a 12-hour light-dark cycle. Cage cleaning was kept to a maximum of once per week and clean/dry nesting material was transferred to new cages during cage change to maintain familiar odours. Cage cleaning was maintained during experiments, although cages were additionally changed following the initiation of antibiotic treatment to prevent coprophagy of microbially-replete faecal pellets.

C57BL/6J GF mice were bred and housed in sterile isolators prior to microbial colonisation or infection experiments. These mice were maintained in positive-pressure isolators and screened every week for contamination using faecal culture, microscopy, and 16S rRNA gene PCR. Consumables were autoclaved at 121°C before introduction into the isolators. For experiments, cages were removed from isolators and opened in a vapourised hydrogen peroxide-sterilised class II safety cabinet. Mice were only handled under aseptic conditions. Mice were then maintained in sterilised bioexclusion cages (IsoCage P, Tecniplast) under positive pressure to prevent external contamination. Mice of both sexes aged between six and nine weeks were used in GF infection studies. GF mouse experiments were performed in a Containment Level 3 facility.

### Bacterial strains

Selected isolates of the MCC were administered to GF mice to screen for resistance to *S.*Tm infection. The details of these isolates are included in Table 1. Non-MCC isolates were also included in this screen, namely *Enterococcus_B lactis* (formerly *Enterococcus faecium*) strain Com15 and *Clostridium_A leptum*.

*S.*Tm strain 14028 (strain designation: CDC 6516-60) was purchased from the American Type Culture Collection (ATCC) and was the main strain used in the screening protocol. Strains SL1344 and M525 were also trialled in pilot studies of the infection protocol. All isolates were stored at –80°C in 25% glycerol for medium term storage.

### Administration of antibiotics via oral gavage

A broad-spectrum antibiotic cocktail of ampicillin (20 mg/mL), vancomycin (10 mg/mL), metronidazole (10 mg/mL), neomycin (20 mg/mL), and ciprofloxacin (3 mg/mL) was prepared using autoclaved drinking water and by stirring and heating to a maximum of 42°C. Antibiotic solutions were sterile filtered (0.22 um filter) prior to administration to mice.

To perform oral gavage, mice were anaesthetised with isofluorane before receiving 200 uL of antibiotic solution intragastrically via a metal ball-tipped feeding needle. Oral gavage was performed each morning daily for seven days using fresh preparations of the antibiotic cocktail.

### Administration of antibiotics in the drinking water

Ampicillin (1 g/L), vancomycin (0.5 g/L), metronidazole (0.5 g/L), and neomycin (1 g/L) were dissolved in autoclaved drinking water using stirring and heating to a maximum of 42°C. Antibiotic solutions were sterile filtered using a 0.22 um vacuum filtration system before being administered to mice in black water bottles. Mice were kept on antibiotic drinking water for 14 days, with antibiotic solutions being renewed each fifth day.

Mice were weighed prior to the initiation of antibiotics and then daily during antibiotic administration; any mouse that was found to lose 20% of their pre-antibiotic starting weight was culled in accordance with the humane endpoints of the study.

### Commensal isolate culture and administration to mice

To culture commensal isolates, cryostocks were removed from the –80°C freezer and kept on dry ice. Stocks were then scraped using a sterile bacterial loop and the scrapings transferred to a sterile Eppendorf tube on dry ice to minimise exposure to oxygen.

Eppendorf tubes were transferred into an anaerobic chamber and scrapings allowed to thaw. Thawed stock cultures were streaked onto YCFA agar plates^56, 57^ and incubated at 37°C until single colonies could be picked – this took between 24 to 96 hours depending on the strain. Single colonies were picked and used to inoculate 20 mL fresh YCFA or MRS broth. Broth was incubated at 37°C until turbid. MRS broth was preferred for culturing *Lactobacillus* species, while YCFA was used for all other species.

For administration to mice, cultures were centrifuged for 10 minutes at 3900 rpm, the supernatant discarded, and cell pellets resuspended in 3 mL of pre-reduced sterile PBS. Commensal culture was administered 200 uL/mouse via oral gavage. Serial dilutions of inoculating cultures were performed pre-and post-gavage to determine inoculum titres, and 16S rRNA gene PCR and sequencing was performed on colonies to verify the identity of the administered bacterial species.

### *Salmonella* Typhimurium culture and mouse infection

For *S.*Tm culture, cryopreserved stocks were scraped using a sterile bacterial loop and transferred to a sterile Eppendorf tube. *S.*Tm was then plated on LB agar plates and incubated at 37°C for 24 hours under aerobic conditions. A single colony was used to inoculate 10 mL of fresh LB broth which was incubated at 37°C overnight shaking at 200 rpm. 100 uL of overnight culture was used to inoculate 3 mL of LB broth (1:30 dilution) which was then incubated at 37°C shaking at 200 rpm for 3 hours to reach log-phase growth and a density of 10^9^ colony-forming units (CFU)/mL. The 3-hour culture was centrifuged at 3,900 rpm for 5 minutes and the supernatant discarded to remove free endotoxin and secreted metabolites. The pellet was then resuspended in 3 mL of cold sterile PBS.

For infections, the washed *S.*Tm culture was further diluted in cold sterile PBS to obtain 10^7^ CFU/mL (1:100 dilution) for SPF experiments or 10^3^ CFU/mL (1:1,000,000 dilution) for GF experiments. 100 uL of the *S.*Tm inoculum was intragastrically administered to each mouse by oral gavage. All *S.*Tm cultures and inocula were plated via serial dilution to confirm that the expected infectious dose had been given.

Mice were weighed on the day of infection, prior to *S.*Tm oral gavage, and then each morning following infection. Any animal that was found to have lost 20% of their pre-infection starting weight was culled in accordance with the humane endpoints of the study. Physical manifestations of severe disease, i.e., hunching, piloerection, glazed eyes, were also considered as humane endpoints, and any mouse exhibiting these symptoms was culled.

### Faecal sample plating and enumeration of faecal titres

Faecal samples were plated to enumerate commensal or pathogen titres, as well as to confirm colonisation and ensure the absence of contamination following monocolonisation of GF mice. Fresh faeces were collected from mice into pre-weighed sterile Eppendorf tubes using aseptic technique. Eppendorf tubes containing faeces were reweighed and the faecal weights recorded. 500 uL of sterile PBS was then added and faecal pellets homogenised using a sterile P1000 pipette tip before eight 1:10 serial dilutions were performed for each sample. For each sample, 10 uL of each dilution was plated in triplicate on pre-reduced and pre-warmed agar plates.

For the SPF mouse infection experiments, faecal samples were taken from mice using aseptic technique following cessation of antibiotic treatment, prior to infection, and then each day from 2 days post infection (dpi). Pre-infection samples were plated on YCFA plates under anaerobic conditions to quantify commensal titres; post infection samples were plated on Salmonella Shigella agar (SSA; ThermoFisher Scientific, CM0099) to enable quantification of *Salmonella* titres. Faeces from 1 dpi were not plated.

For GF mouse experiments, faeces were taken one week after commensal gavage during cage change and then again at two weeks after commensal gavage – the same day as pathogen infection. Samples were plated on YCFA to confirm colonisation and the absence of contamination. Representative colonies for each morphology were picked and processed for 16S rRNA gene sequencing to confirm the identity of the colonising organism.

### 16S rRNA gene PCR and sequencing

Bacterial colonies were picked using a bacterial loop and mixed into a screw cap tube containing DNA-free acid-washed glass beads and 500 uL of sterile PBS. The bead tube was then shaken at speed 6,000 for 30 seconds using a FastPrep Instrument (MPBio). Bead tubes were then centrifuged at 13,200 rpm for 10 minutes to pellet the cell debris, and the supernatant was taken for the PCR reaction.

GoTaq DNA polymerase kits (Promega) were used for PCR reactions. The 16S rRNA gene was amplified using the broad-range bacterial primers Bact-7F (5’-AGA GTT TGA TYM TGG CTC AG-3’) and Bact-1510R (5’-ACG GYT ACC TTG TTA CGA CTT-3’) which amplify the entire length of the 16S rRNA gene resulting in a PCR product of around 1,500 bp. PCR was performed in 50 uL reactions on a T100 thermal Cycler (Bio-Rad) for 30 cycles, according to manufacturer’s instructions. Finally, 10 uL of the PCR product was run on a 1% agarose gel (0.8 g agarose + 80 mL TBE) containing SYBR Safe DNA Gel stain for 15 minutes at 100 V and 200 A. DNA bands were then visualised using UV light. If the reaction had generated an appropriately sized band, the PCR product was submitted to Eurofins Genomics for PCR product clean up and sequencing.

### Enumeration of *Salmonella* titres in mouse tissues following infection

GF mice were monocolonised and infected as described above. Mice were culled at 24 hours post-infection and whole intestines (jejunum to rectum), mesenteric lymph nodes (mLNs), liver, and spleen removed and kept separately in ice-cold PBS. The gallbladder was separated intact from the liver to avoid contamination with its contents.

The remaining mesenteric and peri-intestinal fat was removed. Samples 0.5–1 cm in length were taken from the jejuno-ileal junction, the terminal ileum, the caecum, and the proximal colon for histology; tissues were placed in tissue cassettes and submerged in methacarn (60% methanol, 30% chloroform, 10% glacial acetic acid). Luminal contents were then collected from the caecum into sterile pre-weighed Eppendorf tubes and stored on ice until weighing and plating. Intestines were next separated into the small intestine, caecum, and colon and processed separately. Each tissue was flayed open longitudinally, luminal contents were gently removed, and tissue was washed sequentially in three petri dishes containing ice-cold PBS.

All tissues were then weighed before being manually homogenised in 0.1% Triton X-100 solution. Livers were added to 6-well plates containing 3 mL 0.1% Triton X-100 solution, while mLNs, spleens, and caeca were added to Eppendorf tubes each containing 500 uL of 0.1% Triton X-100 solution. All tissues were kept on ice throughout this process. Following organ homogenisation, serial dilution of the homogenates was performed and plated on SSA plates to quantify *S.*Tm titres. Caecal contents were weighed before being homogenised in 500 uL of sterile PBS. These samples were briefly spun in a microcentrifuge to settle faecal matter and serial dilutions performed on the supernatants. Serial dilutions were plated on SSA plates to quantify *S.*Tm luminal titres.

### Intestinal epithelial and immune cell isolation

Whole intestines and mLNs were removed from mice and transluminal sections taken from the jejuno-ileal junction, terminal ileum, caecum and distal colon for histology. Luminal contents were collected from the terminal ileum, caecum, and distal colon, and kept in pre-weighed Eppendorf tubes on ice until plating or further processing. Peyer’s patches were excised from the small intestine and kept on ice until processing. The intestines were opened longitudinally, cleared of faeces, and then washed three times in cold PBS. To isolate the intraepithelial immune populations, the intestines were cut into 1 cm pieces, washed in cold PBS, and then incubated in 10 mL of PBS with 1 mM dithiothreitol for 10 minutes. Tissues were manually disrupted via shaking for 2 minutes and then strained, collecting the supernatant. Tissues were then incubated in 10 mL of PBS with 30 mM EDTA and 10 mM HEPES at 37°C in a shaker at 200 rpm for 10 minutes, before being manually shaken and strained. The supernatants from these two steps were pooled, filtered at 70 um and the filtrate fractionated using a discontinuous Percoll gradient (80%/40%). Epithelial cells were isolated from the surface of the Percoll, and the intraepithelial immune cells isolated from the interface.

To isolate lamina propria immune cells, the intestines were incubated again in 10 mL of PBS with 30 mM EDTA and 10 mM HEPES at 37°C in a shaker at 200 rpm for 10 minutes, before being manually shaken and strained. Intestinal tissues were then finely chopped and digested in complete HBSS-2 (HBSS, 25 mM HEPES, 1 mM sodium pyruvate) containing 0.05 mg/mL Collagenase VIII (Sigma) and 50 mg/mL DNase I (Sigma) for 1 hour at 37°C, shaking at 80 rpm. Samples were mechanically disrupted by pipetting through serological pipettes and filtered at 70 um. The filtrate was fractionated using a discontinuous Percoll gradient (80%/40%). Lamina propria immune cells were isolated from the interface.

Peyer’s patches and mLNs were processed using the same protocol. Remaining lamina propria and fat was removed, and tissues digested in 3 mL complete HBSS-2 containing 0.5 mg/mL Collagenase-D (Sigma) at 37°C for 30 minutes. Tissues were then forced through 70 um filters to yield single cell suspensions. Collagenase was quenched with 2 mL complete HBSS-2. Cells were then pelleted, and the supernatant discarded.

### Flow cytometry

Immune cell populations were characterised by flow cytometry. Two panels were used to quantify lymphocytes and myeloid cells. Non-viable cells were stained using Live/Dead Fixable Aqua after which surface markers were stained using fluorophore-conjugated primary antibodies (Table 2). For nuclear staining, lymphocytes were permeabilised with CytoFix/CytoPerm (BD) and FoxP3 staining performed in Permeabilisation Buffer (eBioscience). Cells were then fixed in 1% paraformaldehyde in preparation for flow cytometric analysis. Sample data were acquired with an Attune NxT flow cytometer coupled with an Attune CytKick Max autosampler. Data were analysed using FlowJo v10 and R.

**Table 2.**
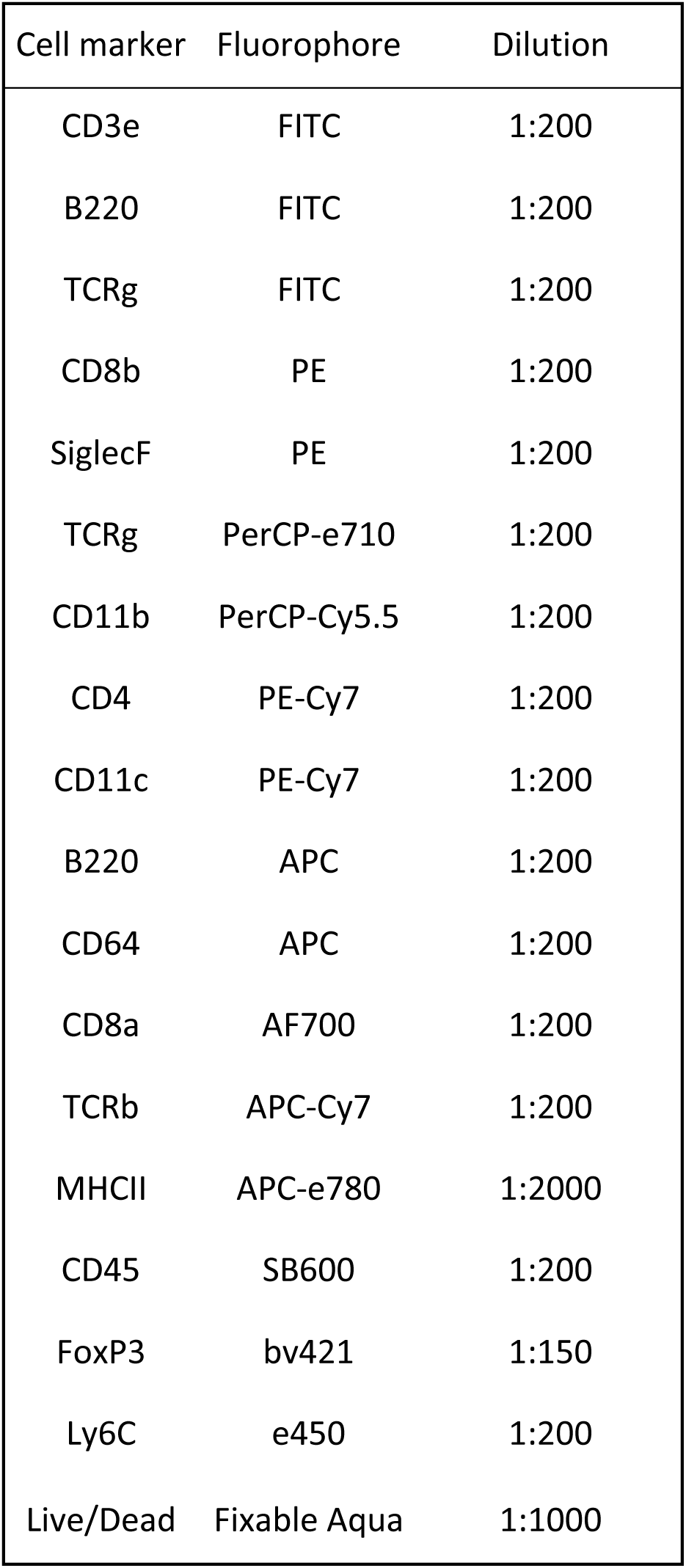
Antibodies for lymphoid and myeloid cell population characterisation by flow cytometry.

### Intestinal histology

For histological analyses, mice were euthanised either prior to infection, on day 14 after commensal monocolonisation, or on day 1 post-infection. Intestinal tissues were excised and samples around 1 cm in length were taken from the jejuno-ileal junction, terminal ileum, the caecum, and the distal colon. Sections containing luminal contents were preferentially taken. Tissue sections were fixed in methacarn (60% methanol, 30% chloroform, 10% glacial acetic acid) for 18 hours at 4°C and then washed three times in 70% ethanol at room temperature for 10 minutes each. Samples were stored in 70% ethanol at 4°C until submission to the histology core facility for paraffin-embedding and sectioning (5 um) and staining with haematoxylin and eosin (H&E) for histology scoring. Additional sections/slides were produced for downstream immunofluorescence (IF) or fluorescence *in situ* hybridisation (FISH).

### Intestinal pathology scoring

Pathohistological scoring of tissue sections was performed according to a previously published protocol^26^. Histology slides of colon and caecum cross-sections were scored across five domains by two independent and blinded researchers. The domains scored were (1) degree of submucosal oedema, (2) severity of leukocyte infiltration in lamina propria, (3) epithelial integrity, (4) presence of cryptitis and crypt abscesses, (5) the number of polymorphonuclear granulocytes (PMN) infiltrating the lamina propria, and (6) the number of goblet cells. Submucosal oedema was assessed using low-power fields (25× magnification), one for each section. Other domains were assessed across ten high-power fields (400× magnification) taken from different regions of each section.

**Submucosal oedema**

0 — no pathological changes
1 — detectable oedema (submucosal oedema, <50% wall thickness)
2 — moderate oedema (submucosal oedema, 50 to 80% wall thickness)
3 — profound oedema (submucosal oedema, ≥80% wall thickness).

**Mononuclear (mixed leukocyte) infiltrate**

0 — none; normal number of mononuclear cells
1 — low; slightly more than normal infiltration
2 — moderate; many cells present and some evidence of tissue distension
3 — high; extreme infiltration, often with tissue architecture disruption.

**Epithelial integrity**

0 — no pathological changes detectable
1 — epithelial desquamation
2 — erosion of the epithelial surface (gaps of 1 to 10 epithelial cells per lesion)
3 — epithelial ulceration (gaps of >10 epithelial cells per lesion, or granulation tissue below the epithelium).

**Cryptitis and crypt abscesses**

0 — none; no neutrophils between epithelial cells or in crypts
1 — low; some evidence of neutrophil infiltration of the crypt epithelium
2 — moderate; many neutrophils between crypt epithelial cells, some in the crypts
3 — high; abundant crypt abscesses (neutrophils in the crypts).

**PMN infiltration into the lamina propria**

0 — <5 PMN/high-power field
1 — 5 to 20 PMN/high-power field
2 — 21 to 60 PMN/high-power field
3 — 61 to 100 PMN/high-power field
4 — >100 PMN/high-power field.

N.B. Where transmigration of PMN into the intestinal lumen was observed, sections were given a score of at least 3.

**Goblet cells**

0 — >28 goblet cells/high-power
1 — 11 to 28 goblet cells/high-power field
2 — 1 to 10 goblet cells/high-power field
3 — <1 goblet cell/high-power field.

For each sample, the two independent scores across all sections were averaged for each domain and the combined pathological score for each sample was determined as the sum of these averaged scores.

### Fluorescence *in situ* hybridisation

FISH was performed using a published protocol^58, 59^ and the universal bacterial FISH primer, Eub338 (5’-GCT GCC TCC CGT AGG AGT-3’), that was conjugated to FITC at both termini.

Fresh hybridisation buffer (0.9 M NaCl, 20 mM Tris-HCl (pH 7.2), 0.1% SDS) was warmed to 50°C and used to dilute the bacterial DNA probe 1:300. 50 uL/slide of staining solution was added directly onto deparaffinised tissue slides and then gently covered with a piece of parafilm cut to the size of a coverslip. Slides were incubated in a humidified box shielded from light for 3 hours at 50°C. Tissues were then washed twice in warm wash buffer (0.9 M NaCl, 20 mM Tris-HCl (pH 7.2)) for 10 minutes each. Next, Hoescht stain was diluted 1:3000 in wash buffer and 100 uL added directly to slides. Slides were incubated for 5 minutes at room temperature while shielded from light before being washed once more for 10 minutes in wash buffer. Slides were then thoroughly dried, one drop of ProLong Diamond Antifade Mountant directly applied to each slide and a cover glass carefully positioned. Slides were kept at 4°C overnight prior to imaging.

### Immunofluorescence

IF was performed on post-*S.*Tm infection samples to visualise the interaction of *S.*Tm with the intestinal epithelium. *S.*Tm was stained using rabbit antiserum against the O4 antigen (TR1302, Sifin), intestinal mucus was stained using mouse anti-Mucin 2/MUC2 antibody (ccp58; sc-7314, Santa Cruz), and phalloidin-AF488 (ab176753, Abcam) and DAPI (D1306, ThermoFisher Scientific) were used to visualise the actin cytoskeleton and host nuclei respectively.

To perform IF, slides were deparaffinised and kept in PBS for 30 minutes. Antigen retrieval was performed using heat-induced epitope retrieval with a sodium citrate buffer (10 mM Sodium citrate, 0.05% Tween 20, pH 6.0). The buffer was then heated to 95°C in a water bath and the slides incubated at this temperature for 20 minutes. The slides were allowed to cool to room temperature for 30 minutes after which they were washed in MilliQ water for 5 minutes and then PBS for 10 minutes. In preparation for antibody staining, slides were incubated in blocking buffer (2% BSA, 0.5% Triton, in PBS) for 20 minutes at room temperature. Primary antibodies were diluted in blocking buffer (rabbit anti-O4, 1:10 dilution; mouse anti-MUC2, 1:50 dilution) to make a primary staining master-mix. 50 uL primary antibody solution was added directly to slides and left to stain overnight at 4°C. Slides were then washed three times in PBS for 5 minutes each. Secondary antibodies (anti-rabbit-AF568 (A-11011, Invitrogen), anti-mouse-AF647 (ab150115, Abcam)) were diluted 1:1000 in blocking buffer and phalloidin-AF488 conjugate was added to yield a final concentration of 1X. 50 uL of secondary antibody and phalloidin staining mixture was added directly to slides and incubated at room temperature for 2 hours. Slides were then washed three times in PBS for 5 minutes each. Finally, 50 uL of 300 nM DAPI solution (diluted in PBS) was added to each slide and incubated for 5 minutes. Slides were washed three times in PBS for 5 minutes each and then dried thoroughly. One drop of ProLong Diamond Antifade mounting medium was directly applied to each sample and a cover glass carefully positioned to prevent bubble formation. Slides were kept at 4°C overnight prior to imaging.

### Confocal microscopy and image analysis

FISH and IF stained slides were visualised using a Leica TCS SP8 Lightning confocal laser scanning microscope (Leica Microsystems). For figure production, images were processed using Fiji-2. Auto-adjustment of brightness and contrast was used to optimise colour handling in captured images.

### Scoring mucus penetration by commensal microbes

Commensal microbe interactions with the intestinal epithelium were qualitatively ascertained using a mucus penetration score. Ten high-power fields were taken of each section and scored by an independent, blinded researcher according to the below rubric:

0 — bacteria are completely excluded from the inner mucus layer
1 — some bacteria can be seen in the inner mucus layer, but there is no contact
2 — many bacteria can be seen in the inner mucus layer with sporadic contact with the epithelium, however there is still clear evidence of a distinct mucus layer
3 — many bacteria can be seen in the inner mucus layer with many contact points between bacteria and the epithelium, there is less evidence of a clear mucus layer.
4 — significant contact of bacteria with the epithelium, with limited evidence of a mucus layer
5 — complete contact of the bacteria with the epithelium, no evidence of a mucus layer.

### RNA extraction from intestinal epithelial cells

IECs were isolated during the first Percoll gradient step during the immune cell isolation protocol. The epithelial cell layer was pipetted from the surface of the Percoll gradient using a P1000 pipette and transferred to a 15 mL Falcon tube containing 8 mL of HBSS-2. IECs were centrifuged at 4°C for 5 minutes at 2,000 rpm and the supernatant discarded. IECs were then resuspended in 1 mL of HBSS-2 and transferred to a PCR-grade 1.5 mL Eppendorf tube. Tubes were centrifuged for at 4°C for 1 minute at 900 x g and supernatant discarded.

The cell pellet was resuspended in 1 mL of TRIzol (Invitrogen), pipetting up and down until it was completely homogenised. The tubes were vortexed briefly before being centrifuged at 4°C for 10 minutes at 12,000 x g to pellet fat and insoluble material. The top soluble fraction was then transferred to a new Eppendorf tube and immediately frozen at –80°C until RNA extraction. RNA was extracted from TRIzol according to the manufacturer’s instructions.

RNA concentration and purity were calculated using a Nanodrop. For each sample, RNA concentrations were diluted to 200 ng/uL in RNase-free water. An A260:A280 ratio over 1.8 was considered high purity. If RNA was low purity, then a clean-up protocol was implemented. For this, RNA sample volumes were made up to 500 uL with RNase-free water and 50 uL of 3M sodium acetate (pH 5.5) was added with 10 ug glycogen, depending on whether it was a low yield sample. 500 uL of isopropanol was then added, the tube mixed well, and then incubated for 20 minutes. RNA was pelleted by spinning at 12,000 x g at 4°C and supernatant removed. The pellet was then washed twice with 300–500 uL ice cold ethanol. Finally, the RNA pellet was left to air dry for 15 minutes before resuspension in RNase-free water.

### RNA quantification by reverse transcription-quantitative polymerase chain reaction

Sequences for each gene of interest was accessed via UniProt and then analysed as a PCR template using the NCBI Primer-BLAST web interface tool^60^. Primers were designed to produce a PCR product with an amplicon size of 70–200 bp and were checked for specificity against *Mus musculus* RefSeq mRNA database. Results for potential primers were then manually screened and the two most optimal primer pairs chosen that minimised predicted self-complementarity. The sequences for primers used in this study are provided in Table 3.

**Table 3.**
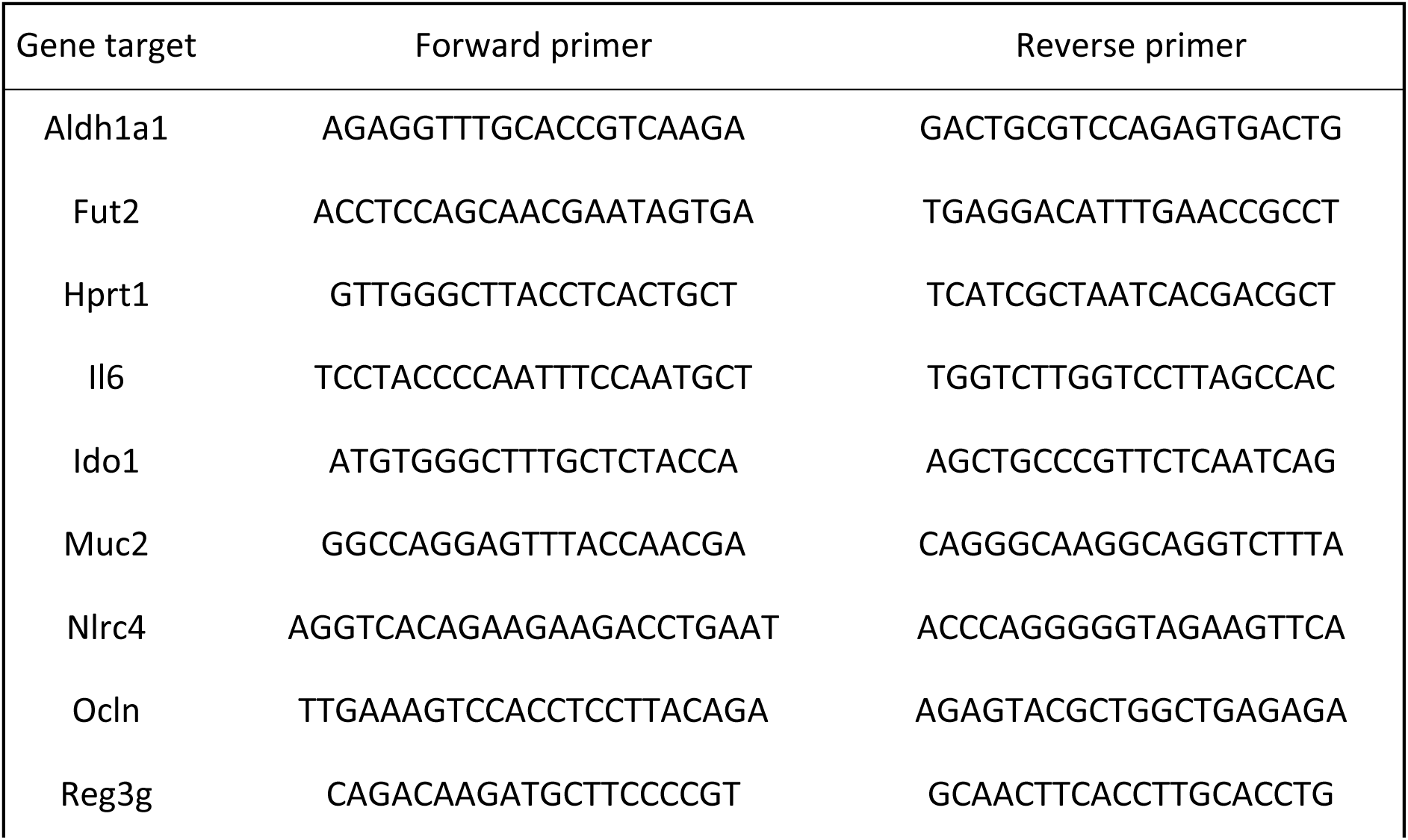
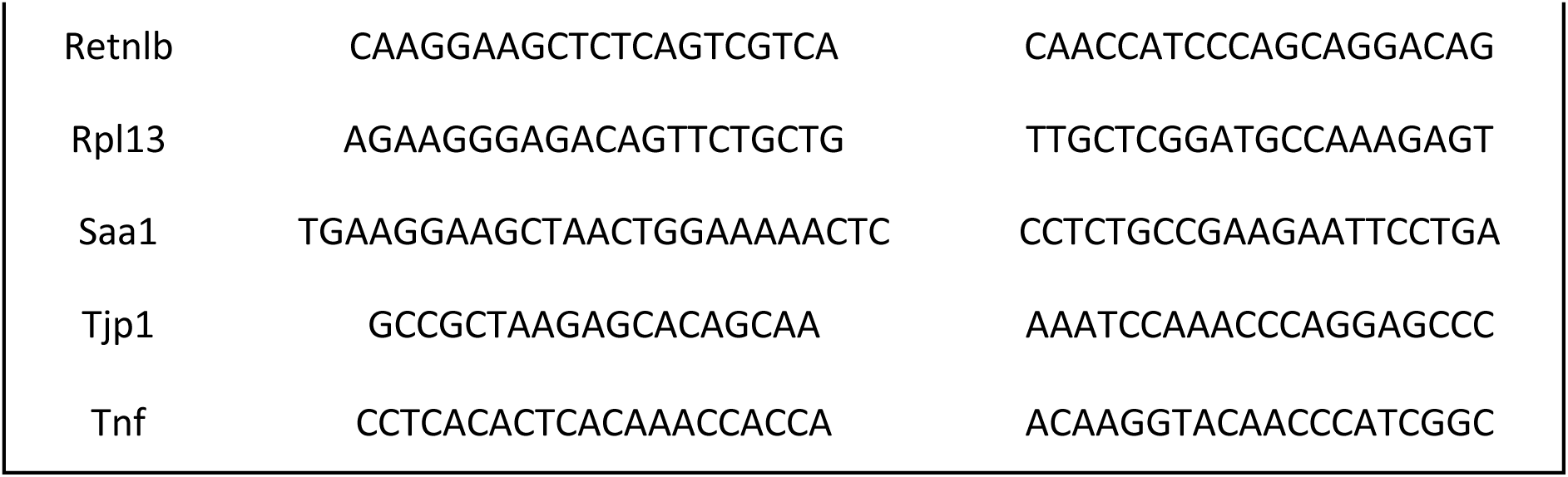
qPCR primers for IEC gene expression. All primer sequences are provided for the 5’– 3’ direction.

To perform RTqPCR, cDNA was first synthesised from the extracted RNA using the Bio-Rad iScript kit and a T100 Thermal Cycler (Bio-Rad) as per the manufacturer’s instructions. Resulting cDNA was diluted to 5 ng/uL in nuclease-free water prior to qPCR. qPCR reactions were performed in 384-well plates using the iTaq Universal SYBR Green Supermix (Bio-Rad) as per manufacturer’s instructions on a QuantStudio 7 Flex qPCR machine (‘fast’ mode, 40 cycles). Data were exported and analysed in R. All runs were quality controlled using melt curve profiles. Low-quality runs, with unexpected melt curve patterns including bimodal melt curves and different peak melt temperatures from reference, were excluded from analyses.

The mean Ct value for two housekeeping genes, *rpl13* and *hprt1*, was calculated per sample and the relative expression of test genes calculated using the formula

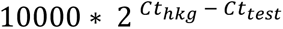

where Ct_hkg_ is the mean Ct value of the housekeeping genes and Ct_test_ is the Ct value of the target gene of interest. Relative expression levels of genes were then compared between treatment groups using t-tests.

### *Ex vivo* faecal spiking assay

Fresh faecal pellets were collected from GF and gnotobiotic mice monocolonised with either *E. clostridioformis* A92, *E. coli* A7, or *A. faecis* A14 into pre-weighed Eppendorf tubes. Tubes containing luminal contents were then reweighed and rapidly transferred to an anaerobic cabinet. Pre-reduced sterile PBS was added to each sample such that faecal pellets were diluted to 0.5 g/mL. Samples were homogenised and 200 uL of the faecal slurry was taken to perform the faecal spiking assay.

For faecal spiking assays, a preparation of 10^6^ CFU/mL *S.*Tm was prepared under aerobic conditions as described above. 20 uL was added to homogenised samples such that the final *S.*Tm titre was 10^5^ CFU/mL. Spiked faecal samples were then incubated under anaerobic conditions at 37°C for 24 hours. To quantify *S.*Tm titres throughout the experiment, serial dilutions were performed at 4, 8, and 24 hour timepoints using 10 uL of each sample. Serial dilutions were plated on preheated SSA plates and then incubated under aerobic conditions at 37°C for 24 hours until colonies were quantified.

### *In vitro* colonisation resistance assays

*S.*Tm growth was tracked using CFU quantification in commensal isolate stationary phase broth culture and sterile-filtered spent media, while OD was used as a proxy for growth in the spent media glucose rescue assays.

For all protocols, commensal isolates were grown on agar as described above and single colonies inoculated into pre-reduced LB broth which was then incubated at 37°C under anaerobic conditions for 36 hours to reach stationary phase. Growth curves were generated for all isolates using CFU/mL to confirm that isolates grew in LB and determine the time required to reach stationary phase. For the spent media assays, stationary phase media was centrifuged at 3,900 rpm for 10 minutes and the supernatant sterile filtered using 22 um filters into sterile 50 mL tubes. Stationary phase media and spent media were each split into three 10 mL aliquots. For the glucose rescue assays, spent media was split into 2 mL aliquots and sterile 10% glucose solution added to aliquots to make up final glucose concentrations of 0%, 0.02%, 0.05%, 0.1%, and 0.2%. Following this, 200 uL of each concentration was added in triplicate to a flat-bottom 96-well plate.

For *S.*Tm inoculation, a fresh *S.*Tm preparation of 10^8^ CFU/mL was added 1:100 to each aliquot or well. For stationary phase and spent media experiments, inoculated media was then incubated without shaking under anaerobic conditions at 37°C for 24 hours. At 4-, 8-, and 24-hour timepoints, tubes were gently swirled and 100 uL of each culture was taken and serial dilutions performed. Serial dilutions were plated on SSA (stationary) or LB (spent) plates to quantify *S.*Tm CFU titres. For glucose rescue assays, *S.*Tm was directly inoculated into wells of the 96-well plate. The OD of each well was then measured after 24 hours of stationary incubation at 37°C under anaerobic conditions.

## References

1. Balasubramanian, R. et al. The global burden and epidemiology of invasive non-typhoidal Salmonella infections. Hum. Vaccines Immunother. 15, 1421–1426 (2019).

2. Majowicz, S. E. et al. The Global Burden of Nontyphoidal Salmonella Gastroenteritis. Clin. Infect. Dis. 50, 882–889 (2010).

3. Kirk, M. D. et al. World Health Organization Estimates of the Global and Regional Disease Burden of 22 Foodborne Bacterial, Protozoal, and Viral Diseases, 2010: A Data Synthesis. PLOS Med. 12, e1001921 (2015).

4. Bernad-Roche, M., Casanova-Higes, A., Marín-Alcalá, C. M., Cebollada-Solanas, A. & Mainar-Jaime, R. C. Salmonella Infection in Nursery Piglets and Its Role in the Spread of Salmonellosis to Further Production Periods. Pathogens 10, 123 (2021).

5. Castro-Vargas, R. E., Herrera-Sánchez, M. P., Rodríguez-Hernández, R. & Rondón-Barragán, I. S. Antibiotic resistance in Salmonella spp. isolated from poultry: A global overview. *Vet*. World 13, 2070–2084 (2020).

6. Tasmin, R., Gulig, P. A. & Parveen, S. Detection of Virulence Plasmid-Encoded Genes in Salmonella Typhimurium and Salmonella Kentucky Isolates Recovered from Commercially Processed Chicken Carcasses. J. Food Prot. 82, 1364–1368 (2019).

7. Chen, Y., Liu, H., Chen, M., Sun, H.-Y. & Wu, Y.-N. The human health burden of non-typhoidal Salmonella enterica and Vibrio parahaemolyticus foodborne gastroenteritis in Shanghai, east China. PloS One 15, e0242156 (2020).

8. Bohnhoff, M., Drake, B. L. & Miller, C. P. Effect of Streptomycin on Susceptibility of Intestinal Tract to Experimental Salmonella Infection. Proc. Soc. Exp. Biol. Med. 86, 132– 137 (1954).

9. Brugiroux, S. et al. Genome-guided design of a defined mouse microbiota that confers colonization resistance against *Salmonella enterica* serovar Typhimurium. Nat. Microbiol. 2, 16215 (2017).

10. Deriu, E. et al. Probiotic Bacteria Reduce Salmonella Typhimurium Intestinal Colonization by Competing for Iron. Cell Host Microbe 14, 26–37 (2013).

11. Herp, S. et al. Mucispirillum schaedleri Antagonizes Salmonella Virulence to Protect Mice against Colitis. Cell Host Microbe 25, 681–694.e8 (2019).

12. Kamada, N. et al. Regulated Virulence Controls the Ability of a Pathogen to Compete with the Gut Microbiota. Science 336, 1325–1329 (2012).

13. Lam, L. H. & Monack, D. M. Intraspecies competition for niches in the distal gut dictate transmission during persistent Salmonella infection. PLoS Pathog. 10, e1004527 (2014).

14. Liou, M. J. et al. Host cells subdivide nutrient niches into discrete biogeographical microhabitats for gut microbes. Cell Host Microbe 30, 836–847.e6 (2022).

15. Litvak, Y. et al. Commensal Enterobacteriaceae Protect against Salmonella Colonization through Oxygen Competition. Cell Host Microbe 25, 128–139.e5 (2019).

16. Maier, L. et al. Microbiota-derived hydrogen fuels Salmonella typhimurium invasion of the gut ecosystem. Cell Host Microbe 14, 641–651 (2013).

17. Nguyen, B. D. et al. Import of Aspartate and Malate by DcuABC Drives H2/Fumarate Respiration to Promote Initial Salmonella Gut-Lumen Colonization in Mice. Cell Host Microbe 27, 922–936.e6 (2020).

18. Velazquez, E. M. et al. Endogenous Enterobacteriaceae underlie variation in susceptibility to Salmonella infection. Nat. Microbiol. 4, 1057 (2019).

19. Jacobson, A. et al. A Gut Commensal-Produced Metabolite Mediates Colonization Resistance to Salmonella Infection. Cell Host Microbe 24, 296–307.e7 (2018).

20. Sorbara, M. T. et al. Inhibiting antibiotic-resistant Enterobacteriaceae by microbiota-mediated intracellular acidification. J. Exp. Med. 216, 84–98 (2019).

21. Byndloss, M. X. et al. Microbiota-activated PPAR-γ signaling inhibits dysbiotic Enterobacteriaceae expansion. Science 357, 570–575 (2017).

22. Cash, H. L., Whitham, C. V., Behrendt, C. L. & Hooper, L. V. Symbiotic Bacteria Direct Expression of an Intestinal Bactericidal Lectin. Science 313, 1126–1130 (2006).

23. Vaishnava, S. et al. The antibacterial lectin RegIIIgamma promotes the spatial segregation of microbiota and host in the intestine. Science 334, 255–258 (2011).

24. Tsolis, R. M. et al. Of mice, calves, and men. Comparison of the mouse typhoid model with other Salmonella infections. Adv. Exp. Med. Biol. 473, 261–274 (1999).

25. Tsolis, R. M., Xavier, M. N., Santos, R. L. & Bäumler, A. J. How to become a top model: impact of animal experimentation on human Salmonella disease research. Infect. Immun. 79, 1806–1814 (2011).

26. Barthel, M. et al. Pretreatment of Mice with Streptomycin Provides a Salmonella enterica Serovar Typhimurium Colitis Model That Allows Analysis of Both Pathogen and Host. Infect. Immun. 71, 2839–2858 (2003).

27. Bohnhoff, M. & Miller, C. P. Enhanced susceptibility to Salmonella infection in streptomycin-treated mice. J. Infect. Dis. 111, 117–127 (1962).

28. Gantois, I. et al. Butyrate specifically down-regulates salmonella pathogenicity island 1 gene expression. Appl. Environ. Microbiol. 72, 946–949 (2006).

29. Hung, C.-C. et al. The intestinal fatty acid propionate inhibits Salmonella invasion through the post-translational control of HilD. Mol. Microbiol. 87, 1045–1060 (2013).

30. Kohli, N. et al. The microbiota metabolite indole inhibits Salmonella virulence: Involvement of the PhoPQ two-component system. PLoS ONE 13, (2018).

31. Zhang, Z. J., Pedicord, V. A., Peng, T. & Hang, H. C. Site-specific acylation of a bacterial virulence regulator attenuates infection. Nat. Chem. Biol. 16, 95–103 (2020).

32. Thiemann, S. et al. Enhancement of IFNγ Production by Distinct Commensals Ameliorates Salmonella-Induced Disease. Cell Host Microbe 21, 682–694.e5 (2017).

33. Ivanov, I. I. & Littman, D. R. Segmented filamentous bacteria take the stage. Mucosal Immunol. 3, 209–212 (2010).

34. Ivanov, I. I. et al. Induction of Intestinal Th17 Cells by Segmented Filamentous Bacteria. Cell 139, 485–498 (2009).

35. Pedicord, V. A. et al. Exploiting a host-commensal interaction to promote intestinal barrier function and enteric pathogen tolerance. Sci. Immunol. 1, (2016).

36. Rangan, K. J. et al. A secreted bacterial peptidoglycan hydrolase enhances tolerance to enteric pathogens. Science 353, 1434–1437 (2016).

37. Beresford-Jones, B. S. et al. The Mouse Gastrointestinal Bacteria Catalogue enables translation between the mouse and human gut microbiotas via functional mapping. Cell Host Microbe (2021) doi:10.1016/j.chom.2021.12.003.

38. Stecher, B. Establishing causality in Salmonella-microbiota-host interaction: The use of gnotobiotic mouse models and synthetic microbial communities. Int. J. Med. Microbiol. 311, 151484 (2021).

39. Rivera-Chávez, F. et al. Depletion of butyrate-producing Clostridia from the gut microbiota drives an aerobic luminal expansion of Salmonella. Cell Host Microbe 19, 443–454 (2016).

40. Eberl, C. et al. E. coli enhance colonization resistance against Salmonella Typhimurium by competing for galactitol, a context-dependent limiting carbon source. Cell Host Microbe 29, 1680–1692.e7 (2021).

41. Reeves, A. E., Koenigsknecht, M. J., Bergin, I. L. & Young, V. B. Suppression of Clostridium difficile in the Gastrointestinal Tracts of Germfree Mice Inoculated with a Murine Isolate from the Family Lachnospiraceae. Infect. Immun. 80, 3786–3794 (2012).

42. Forster, S. C. et al. Identification of gut microbial species linked with disease variability in a widely used mouse model of colitis. Nat. Microbiol. 7, 590–599 (2022).

43. Zmora, N. et al. Personalized Gut Mucosal Colonization Resistance to Empiric Probiotics Is Associated with Unique Host and Microbiome Features. Cell 174, 1388–1405.e21 (2018).

44. Sorini, C., Cardoso, R. F., Gagliani, N. & Villablanca, E. J. Commensal Bacteria-Specific CD4+ T Cell Responses in Health and Disease. Front. Immunol. 9, (2018).

45. Johansson, M. E. V. et al. Normalization of host intestinal mucus layers requires long-term microbial colonization. Cell Host Microbe 18, 582–592 (2015).

46. Clay, S. L., Bravo-Blas, A., Wall, D. M., MacLeod, M. K. L. & Milling, S. W. F. Regulatory T cells control the dynamic and site-specific polarization of total CD4 T cells following Salmonella infection. Mucosal Immunol. 13, 946–957 (2020).

47. Round, J. L. & Mazmanian, S. K. Inducible Foxp3+ regulatory T-cell development by a commensal bacterium of the intestinal microbiota. Proc. Natl. Acad. Sci. U. S. A. 107, 12204–12209 (2010).

48. Atarashi, K. et al. Induction of Colonic Regulatory T Cells by Indigenous Clostridium Species. Science 331, 337–341 (2011).

49. Geuking, M. B. et al. Intestinal bacterial colonization induces mutualistic regulatory T cell responses. Immunity 34, 794–806 (2011).

50. Johanns, T. M., Ertelt, J. M., Rowe, J. H. & Way, S. S. Regulatory T Cell Suppressive Potency Dictates the Balance between Bacterial Proliferation and Clearance during Persistent Salmonella Infection. PLoS Pathog. 6, e1001043 (2010).

51. Atarashi, K. et al. T_reg_ induction by a rationally selected mixture of Clostridia strains from the human microbiota. Nature 500, 232 (2013).

52. Furusawa, Y. et al. Commensal microbe-derived butyrate induces the differentiation of colonic regulatory T cells. Nature 504, 446 (2013).

53. Propheter, D. C., Chara, A. L., Harris, T. A., Ruhn, K. A. & Hooper, L. V. Resistin-like molecule β is a bactericidal protein that promotes spatial segregation of the microbiota and the colonic epithelium. Proc. Natl. Acad. Sci. 114, 11027–11033 (2017).

54. Hogan, S. P. et al. Resistin-like molecule β regulates innate colonic function: Barrier integrity and inflammation susceptibility. J. Allergy Clin. Immunol. 118, 257–268 (2006).

55. Bergstrom, K. S. B. et al. Goblet Cell Derived RELM-β Recruits CD4+ T Cells during Infectious Colitis to Promote Protective Intestinal Epithelial Cell Proliferation. PLoS Pathog. 11, e1005108 (2015).

56. Duncan, S. H., Hold, G. L., Harmsen, H. J. M., Stewart, C. S. & Flint, H. J. Growth requirements and fermentation products of Fusobacterium prausnitzii, and a proposal to reclassify it as Faecalibacterium prausnitzii gen. nov., comb. nov. Int. J. Syst. Evol. Microbiol. 52, 2141–2146 (2002).

57. Lopez-Siles, M. et al. Cultured Representatives of Two Major Phylogroups of Human Colonic Faecalibacterium prausnitzii Can Utilize Pectin, Uronic Acids, and Host-Derived Substrates for Growth. Appl. Environ. Microbiol. 78, 420–428 (2012).

58. Salzman, N. H. et al. Analysis of 16S libraries of mouse gastrointestinal microflora reveals a large new group of mouse intestinal bacteriabbThe GenBank accession numbers for the clone sequences reported in this paper can be found in Table 1T1; the accession number for isolate MIB-CB3 is AJ418059. Microbiology 148, 3651–3660 (2002).

59. Swidsinski, A., Weber, J., Loening-Baucke, V., Hale, L. P. & Lochs, H. Spatial Organization and Composition of the Mucosal Flora in Patients with Inflammatory Bowel Disease. J. Clin. Microbiol. 43, 3380–3389 (2005).

60. Ye, J. et al. Primer-BLAST: a tool to design target-specific primers for polymerase chain reaction. BMC Bioinformatics 13, 134 (2012).

